# Effects of CTCF on the regulatory landscape of the mouse *Sox2* locus

**DOI:** 10.64898/2026.03.16.711995

**Authors:** Mathias Eder, Christina J.I. Moene, Marantha Kaagman, Luca Braccioli, Elzo de Wit, Bas van Steensel

**Affiliations:** Division of Molecular Genetics, Netherlands Cancer Institute, Amsterdam, the Netherlands; Oncode Institute, the Netherlands; Division of Gene Regulation, Netherlands Cancer Institute, 1066 CX Amsterdam, the Netherlands

## Abstract

Regulatory elements that control gene transcription are scattered along the genome, yet it is still poorly understood how their precise positioning affects the regulatory landscape. We previously reported a method to relocate a regulatory element to thousands of positions in a genomic locus and probe the position-dependent impact on gene regulation. Here, we apply this approach to systematically query the role and position-dependence of CTCF binding sites (CBSs), which are thought to modulate communication between distal regulatory elements. We focused on the mouse *Sox2* locus in embryonic stem cells, where the *Sox2* gene is activated by a potent distal enhancer. First, we found that a CBS that is naturally located immediately upstream of the *Sox2* promoter endows this promoter with strong orientation-dependent activation, anywhere within the gene–enhancer interval. Second, throughout this interval, insertion of CBSs alone consistently reduces *Sox2* expression in an orientation-dependent manner, suggesting a polarity in the loop extrusion process. Third, two native CBSs located within and downstream of the enhancer subtly help to confine the activation realm of the enhancer. Together, these results illuminate the interplay between CBSs and the detailed regulatory landscape of a genomic locus.

## Introduction

Transcriptional regulation of genes is highly dependent on the interplay between *cis*-regulatory elements including promoters, enhancers, and elements that control genome folding. Tissue– and cell type-specific regulation of gene expression is often conferred by enhancers that can be located tens or hundreds of kilobases (kb) from their target gene (*1*).

The long linear genomic distances can be bridged by looping mechanisms that bring enhancers and promoters in close proximity in three-dimensional space (*1, 2*). One of the key players in this process is CCCTC-binding factor (CTCF) together with the cohesin complex. CTCF stalls loop extrusion by cohesin in an orientation-dependent manner, thus shaping chromatin loops and regulatory neighbourhoods that can constrain enhancer-promoter communication (*3–7*). Some promoters are bound by CTCF to form long-range contacts with enhancers (*8*). Furthermore, CTCF motifs are often enriched at the boundaries of topologically associated domains (TADs), regions of preferential self-interaction in the genome (*9, 10*).

Acute depletion of CTCF leads to rapid loss of many CTCF-anchored loops and weakening of TAD structure genome-wide (*5, 6*). However, the transcriptional consequences of these architectural perturbations are often modest and limited to a subset of genes. Manipulation of individual CTCF binding sites (CBSs) underscores their diverse and often subtle regulatory roles. Deletion of individual CBSs has variable impact on gene regulation, often with no or only minor effects (*11–14*). A few studies reported that ectopic insertion of CBSs can lead to the formation of new, alternative chromatin loops that in some cases “trap” enhancers into smaller, isolated sub-structures (*15–19*). This can be accompanied by a reduction in local gene expression (*15, 19*), although in some cases this effect is very subtle and requires multiple CBS in tandem (*17, 18*). The degree of this interference was also noted to be dependent on the orientation of the CTCF site (*17–20*), supporting the loop extrusion model where CTCF serves as a directional barrier. Besides the effect of CTCF on local E-P loops, CTCF has an important role in restricting ectopic gene activation by enhancers (*21*).

In addition to their frequent occurrence at loop anchors and domain boundaries, CBSs are commonly found near transcription start sites (TSSs), often within a few kilobases upstream of promoters. In some contexts, promoter-proximal CTCF contributes to distal enhancer responsiveness, consistent with a role in organizing promoter connectivity within regulatory neighborhoods (*8*). However, CTCF binding close to the TSS of promoters can also modulate transcription through mechanisms that do not rely on long-range looping, for example by modulating transcription directly (*22*).

Generally, it remains difficult to predict the regulatory impact of individual CBSs, because the context-dependencies are complex and are still poorly understood. Our understanding of this logic will require a detailed knowledge of the regulatory landscape of each locus. Moreover, it is likely that the effects of CBSs on gene activity depend on their precise position relative to the other regulatory elements, but so far this has not been systematically tested. Here we describe a detailed exploration of such position-dependent effects of CBSs, in the context of the entire local landscape of a genomic locus.

We recently developed a method to relocate a regulatory element to thousands of alternative positions across a locus of several hundred kb, by making use of the Sleeping Beauty transposable element. By combining pooled transposition, phenotypic sorting, and mapping of insertion sites, this “hopping” strategy enables the construction of quantitative functional landscapes that reveal how regulatory output depends on genomic position at scale (*23*).

We previously applied this method to investigate the logic of the linear arrangement of a promoter-enhancer pair in the mouse *Sox2* locus. In mouse embryonic stem cells (mESCs) expression of *Sox2* is primarily driven by the *Sox2* control region (SCR), a ∼10 kb cluster of enhancers located 100 kb downstream of the gene (*24, 25*). By large-scale hopping of the *Sox2* promoter linked to a reporter throughout this locus, we found that the activation realm of the SCR is sharply confined within the *Sox2*-SCR interval, and that the *Sox2* gene itself is partially responsible for this confinement (*23*).

Here, we use three complementary sets of hopping experiments to dissect how endogenous and ectopic CBSs shape the *Sox2* regulatory landscape. In each case, we hopped constructs with CBSs, the *Sox2* promoter, or both elements, into thousands of positions throughout the locus and determined the effects on transcriptional activity. First, we test how a CBS that is naturally located upstream of the *Sox2* promoter alters the response of a *Sox2* promoter reporter to the SCR. We found that this CBS confers strong motif-orientation-dependent effects on promoter activation. Second, we investigated how insertion of CBSs (without a promoter) throughout the locus affects the ability of the SCR to activate the endogenous *Sox2* gene. The results show that CBSs insertions anywhere in the *Sox2*-SCR interval cause consistent orientation-dependent reduction of *Sox2* expression. Third, we tested how two native CBSs located near the SCR impact the regulatory landscape of the entire locus. Together, the results provide new insights into the interplay between CBSs and the overall regulatory landscape of a locus, and further illustrate the ability of the hopping approach to quantify position-dependent regulatory logic in native chromatin.

## Results

### Experimental design

To study the effect of different *Sox2* reporters in the native *Sox2* locus, we made use of a previously engineered clonal cell line (*23*) derived from F1-hybrid (129/Sv:CAST/EiJ) mESCs that has both *Sox2* alleles tagged with two different fluorophores: mCherry for the 129 allele and eGFP for the CAST allele **(Fig. 1A)**. Furthermore, the cell line carries a SB transposon with a recombination cassette 6 kb upstream of the endogenous *Sox2* gene, which is 116 kb upstream of the SCR. We refer to this as launch pad (LP) –116 kb. The recombination cassette consists of a double selection marker (HyTK) flanked by a pair of heterotypic Flp recombinase recognition sites (FRT/F3), allowing for easy exchange of the DNA sequence within the transposon **(Fig. 1A)**. This transposon can be efficiently relocated to hundreds of random positions in the locus by transient transfection with a plasmid expressing SB transposase (*23*).

**Fig 1.**
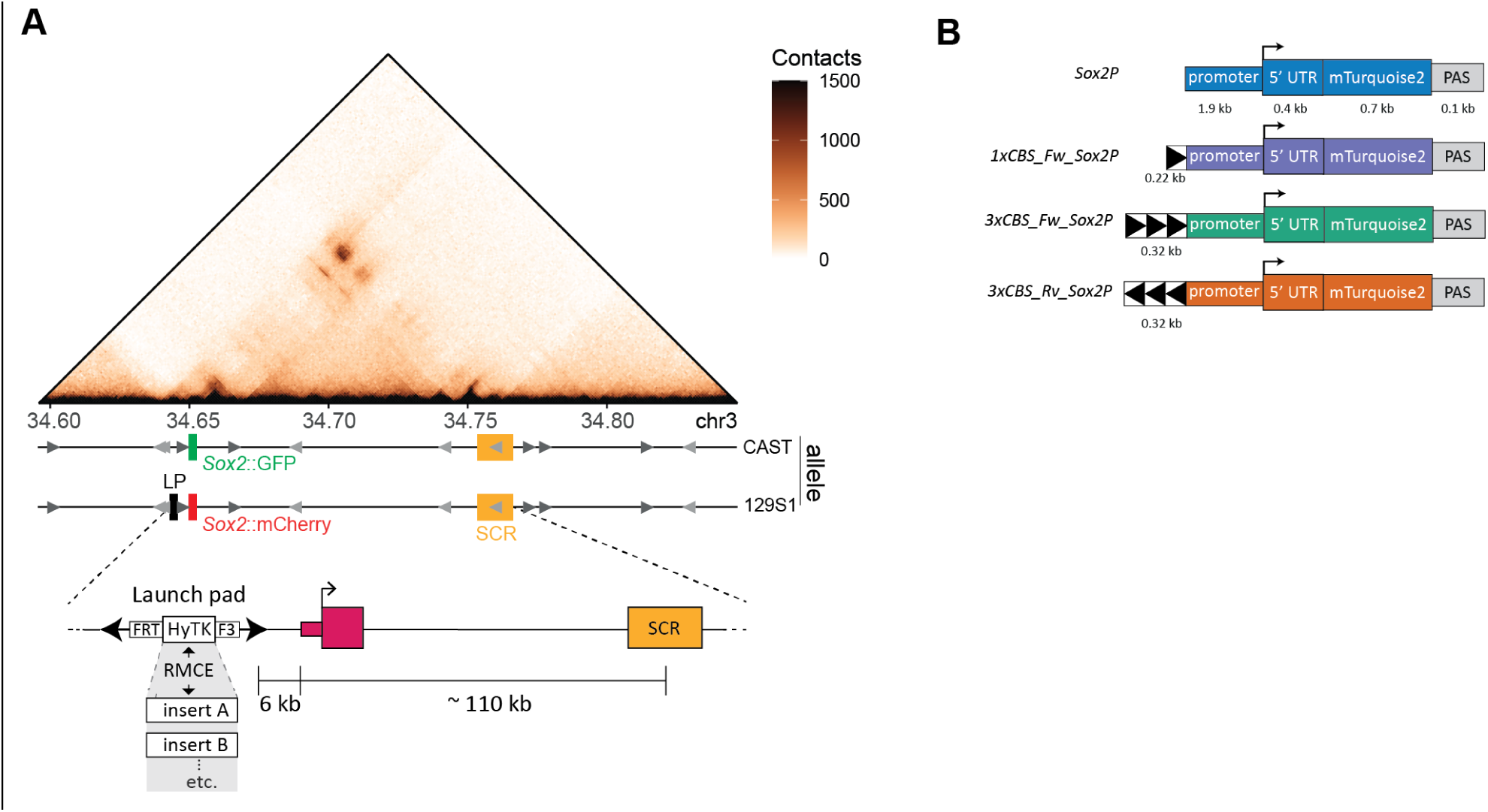
Overview of the murine *Sox2* locus and the different *Sox2* reporter constructs. **(A)** Overview of the *Sox2* locus. (Top to bottom) RCMC contact maps (1-kb resolution) in mESCs from (*26*); positions of SCR, eGFP/mCherry tagged *Sox2* alleles and launch pad (–116 kb from SCR on 129S1 allele) containing SB with a hygromycin phosphotransferase-thymidine kinase (HyTK) recombination cassette flanked by FRT and F3 recombination sites. Position and orientation of endogenous CTCF binding sites depicted as grey triangles **(B)** Schematic representation of different *Sox2* reporters; all mTurquoise2 reporters are driven by *Sox2* promoter and 5’UTR, followed by the SV40 polyadenylation signal (PAS). 1xCBS_Fw_Sox2P reporter carries 221 bp upstream of the *Sox2* promoter containing the *Sox2* promoter proximal CTCF motif (forward). 3xCBS_Fw_Sox2P reporters carry 329 bp containing three strong forward oriented CTCF binding motifs upstream of the *Sox2* promoter. 3xCBS_Rv_Sox2P reporters carry the same three CTCF binding motifs in reverse orientation.

We previously designed a *Sox2* promoter reporter containing 1.9 kb of the *Sox2* promoter (Sox2P) driving the expression of mTurquoise2 (mTurq), a blue fluorescent protein **(Fig. 1B)** (*23*). To test the effect of CBSs upstream of Sox2P, we designed several new constructs. In reporter 1xCBS_Fw_Sox2P, we added the *Sox2*-proximal CBS in forward orientation upstream of the Sox2P reporter **(Fig. 1B)**. To investigate whether any observed effect may depend on the dosage of CBSs, we designed another construct (3xCBS_Fw_Sox2P) that contains a tandem array of three strong CBSs upstream of the Sox2P reporter. This tandem array consists of three tissue-invariant CBS from different genomic regions, each flanked by 30-50 bp of up– and downstream endogenous sequence (see Methods). In their native context, these CBS are strongly bound by Rad21 and CTCF (*16*). The insulating effect of this CBS arrays has been shown previously (*16, 18*). Finally, to test whether any observed effect is dependent on the CBS orientation, we also designed a 3xCBS_Rv_Sox2P reporter, which has the tandem array in reverse orientation **(Fig. 1B)**. By comparison to the original Sox2P reporter lacking a CBS, these three constructs enabled us to study the effects of CBS presence, dosage and orientation on the activity of the reporter driving mTurq expression, and on the endogenous *Sox2* gene that is tagged with mCherry. We recombined each of these reporters into the SB at LP-116 kb.

### Functional maps of the *Sox2* locus generated by large-scale reporter hopping

Next, we set out to generate detailed expression landscapes of the different reporters in the native *Sox2* locus. For this, we induced hopping of each of the reporters from LP-116 kb by transient transfection with SB transposase **(Fig. 2A)**. Seven days later we monitored reporter fluorescence by flow cytometry. Compared to control cells transfected with PiggyBac transposase (mock control which does not mobilize SB), the resulting cell pools showed an increased spread in fluorescence (particularly in the low expression range) **(Fig. 2B)**, indicating that the relocation affected reporter expression. Next, we divided the full range of reporter expression values into six gates (P1-P6) and sorted pools of cells from each gate. After expansion of the cell pools, we mapped the locations of the reporters in each pool, as well as in a pool of unsorted cells (workflow shown in **Fig. S1A**), as described previously (*23*).

**Fig 2.**
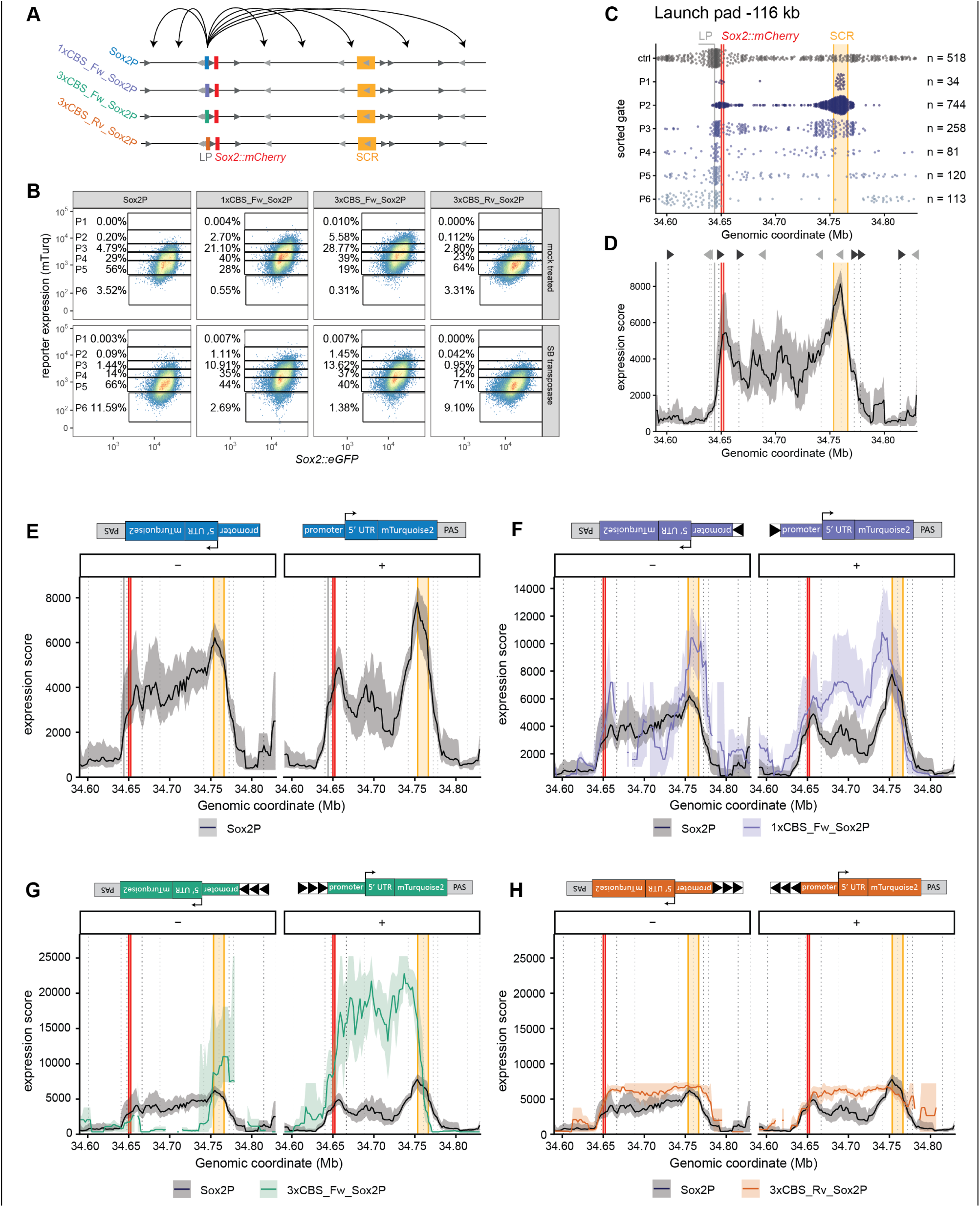
Motif specific effect of CBSs on the activation landscapes of the *Sox2* locus. (**A**) Schematic overview of hopping experiments from LP-116 kb with different reporter constructs. **(B)** Expression levels of different reporters (mTurq) and *Sox2*::eGFP (control) in mock (PiggyBac transfected) and SB-transposase transfected cells, measured by FACS. The percentage of cells in each gate is indicated. Representative result of replicates. **(C)** Mapped integrations from six sorted populations (P1-P6), ranging from highest to lowest mTurq reporter expression (Sox2P) and unsorted control population (ctrl), hopped from launch pad –116 kb. Each dot indicates one mapped integration and n indicates the number of plotted integrations. Combined data from four biological replicates (**Fig. S1B)** (data published previously (*23*)). To allow matched comparisons with the other constructs, we only used published Sox2P data hopped from LP-116 kb. LP: launch pad position; red: *Sox2*::mCherry gene; orange: SCR. **(D)** Expression score of Sox2P integrations mobilized from LP-116 kb, smoothened using a 10 kb window shifted by 1 kb steps and only plotted if at least three integrations per window are present. Shaded region indicates 95% CI. Grey vertical dotted lines and triangles indicate the position and orientation of CBSs. **(E)** Strand specific expression score of Sox2P integrations mobilized from LP-116 kb **(Fig. S2B)**, smoothened using a 20 kb window shifted by 2 kb steps and only plotted if at least three integrations per window are present. Shaded region indicates 95% CI. Grey vertical dotted lines indicate the position and orientation (darkgrey = forward, lightgrey = reverse) of CBSs. **(F-G)** Same as (E) **(F)** 1xCBS_Fw_Sox2P **(G)** 3xCBS_Fw_Sox2P, **(H)** 3xCBS_Rv_Sox2P required at least two integrations per window. Data for F-H consist of two biological replicates combined and y-axis of (G) and (H) are the same for proper comparison.

We performed multiple independent replicates of these experiments for Sox2P (4x), 1xCBS_Fw (2x), 3xCBS_Fw (2x) and 3xCBS_Rv (2x) **(Fig. S1B & S1D)**. Data for the Sox2P experiments from LP-116 kb (**Fig. S1B**) has been published previously (*23*). The integration patterns between replicates were reproducible in the gates with sufficient integrations **(Fig. S1C & S1E)**; therefore, we combined the replicates for further downstream analysis. In total, the analyses below are based on 1350 (Sox2P), 607 (1xCBS_Fw), 405 (3xCBS_Fw) and 721 (3xCBS_Rv) sorted reporter integrations in a 240 kb window centered around the *Sox2* gene and the SCR **(Fig. 2 and Fig. S2)**.

We converted the raw integration data **(Fig. 2C & S2)** into a reporter expression score track (*23*), which is the average expression level in a sliding window across the locus (see figure legend and methods). As reported previously (*23*), the resulting activity landscape of the Sox2P reporter **(Fig. 2C & 2D)** showed strong confinement of reporter activity within the *Sox2*-SCR interval, with local peaks of highest activity around the endogenous *Sox2* gene and the SCR. Reporter activity steeply dropped immediately upstream of *Sox2* and downstream of the SCR **(Fig. 2D)**.

Because the orientation of hopped SBs relative to the launch site is random (*23*), and we anticipated orientation-dependent effects of CTCF, we additionally split the data into two sets according to the + or – orientation of the reporter (we will refer to this as + or – “strand”, to avoid confusion with the orientation of the CBSs relative to the promoter within the reporter constructs), and calculated expression scores separately. Minor activity differences were observed for Sox2P between + and – strand integrations, but these were typically not statistically significant (note the overlapping confidence intervals, **Fig. S2F**), except for slightly higher expression levels of + strand reporters within the SCR.

### CBSs have strand– and orientation-dependent effects on reporter activity

Next, we analysed the hopped CBS-containing reporters. Compared to the Sox2P only reporter, addition of the endogenous CBS to the reporter (1xCBS_Fw_Sox2P) caused notable changes in the reporter activity landscape (**Fig. 2F & S2A**). First, on the + strand, activity increased ∼1.8-fold throughout the *Sox2*-SCR interval. Second, the highest expression of 1xCBS_Fw_Sox2P on the + strand was no longer within the SCR, but between 8 to 18 kb upstream of the SCR. Interestingly, this region displays enhancer like characteristics (**Fig. S2G**), indicated by occupancy of p300 and the transcription factors SOX2, NANOG, and OCT4. This difference is largely accounted for by an increase in P1 integrations in those genomic locations (**Fig. S2C**). In contrast, on the – strand, 1xCBS_Fw_Sox2P showed primarily an increase in activity of reporters within and downstream of the second half of the SCR, while insertions in the *Sox2*-SCR interval showed a more complex pattern, with some areas exhibiting reduced expression compared to Sox2P **(Fig. 2F, S2C & S2G)**.

With 3xCBS_Fw_Sox2P these patterns were greatly augmented (**Fig. 2G & S2A**). On the + strand this reporter showed on average a ∼4.7-fold increase in activity throughout the *Sox2*-SCR interval, compared to Sox2P. The activity in the *Sox2*-SCR interval was higher than within the SCR, with the reporter first reaching the maximum activity again 8 to 18 kb upstream of the SCR. Furthermore, reporter activity on the + strand was more tightly restricted immediately downstream of the SCR (**Fig. 2G**), as indicated by an increased recovery of inactive (P6) reporters in this region (**Fig. S3A & S3B)**. On the – strand, an ∼1.7-fold increased reporter activity was still detected within the SCR and immediately downstream, but throughout the *Sox2*-SCR interval the activity was almost entirely lost **(Fig. 2G & S2H)**. Thus, addition of the potent CBS array to the promoter imposes extreme directionality bias: it further boosts the activity of the reporter on the + strand (except when the reporter is integrated inside or downstream of the SCR) and it strongly reduces reporter activation on the – strand.

Remarkably, when the same 3xCBS array was placed upstream of the reporter promoter but in the reverse orientation (3xCBS_Rv_Sox2P), this directionality bias was completely abolished and the activity landscape was highly similar for the + and – strand integrations. Compared to Sox2P, the activity is modestly increased (1.5-fold for both strands) throughout the *Sox2*-SCR interval, with a monotonous activity landscape without peaks at the SCR or endogenous gene (**Fig. 2H & S2I**). Together, these results demonstrate that CBSs inserted just upstream of the promoter can strongly affect the response to the SCR, most notably when the CBS is in the same orientation as the promoter. These effects are more pronounced with the more potent 3xCBS than with the single endogenous CBS.

### Clonal analysis confirms hopping measurements and suggests some clonal variation

While our hopping strategy provides data for hundreds of integrations, the measurements of individual integrated reporter activities are of limited accuracy due to the rather coarse binning into six gates that span a large expression range. For more precise measurements, we established a series of clonal cell lines with each reporter inserted in two different locations. For this, we chose LP-116 kb (the launch site of the hopping experiments) and a position 39 kb upstream of the SCR (LP –39 kb) **(Fig. 3A)**. In multiple independent clones we precisely measured the reporter mTurq levels by flow cytometry. We then normalized all reporter constructs to the Sox2P control in LP-116 kb and displayed the relative change.

**Fig 3.**
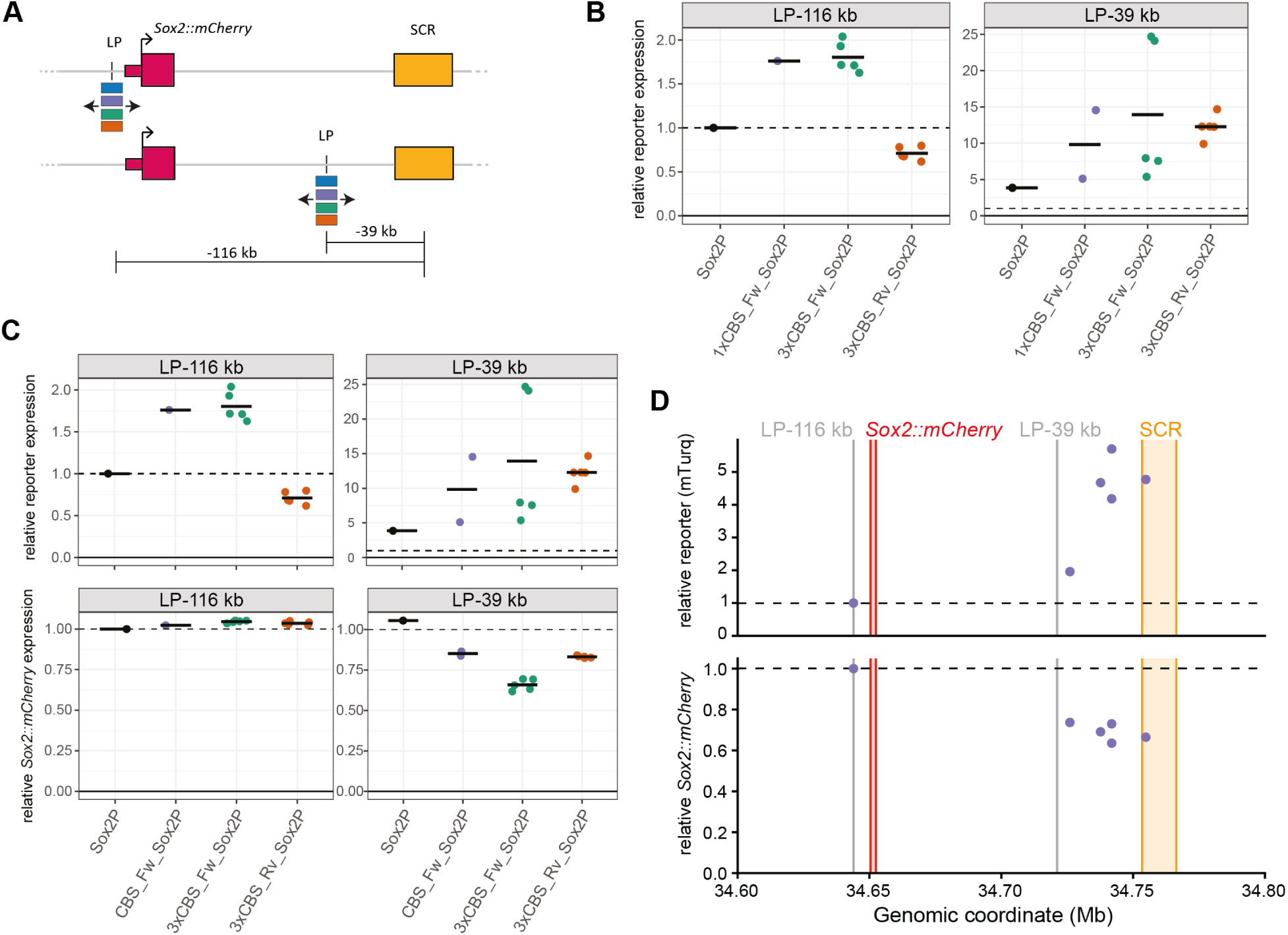
Position and orientation specific effect of CBS on reporter and endogenous *Sox2* expression. (**A**) Schematic representation of testing the different constructs (Sox2P, 1xCBS_Fw_Sox2P, 3xCBS_Fw_Sox2P, 3xCBS_Rv_Sox2P) in two genomic positions, LP-116 kb or LP-39 kb. **(B)** Relative reporter expression (mTurq) of clones containing the Sox2P, 1xCBS_Fw_Sox2P, 3xCBS_Fw_Sox2P or 3xCBS_Rv_Sox2P reporter construct in LP-116 kb or LP-39 kb. Reporter expression was background subtracted, normalized to eGFP and then normalized to the Sox2P reporter expression at LP-116 kb measured on the same day. Each dot represents a clonal cell line. The vertical line indicates the mean reporter expression of the clones. Dashed line indicates the reporter expression of Sox2P at LP-116 kb. **(C)** Top: Same as in (B), Bottom: relative *Sox2*::*mCherry* expression, normalized to the *Sox2*::*mCherry* level of Sox2P in LP-116 kb (C). **(D)** Top: relative reporter expression of clones containing the 1xCBS_Fw_Sox2P construct. Reporter expression was background subtracted, normalized to eGFP and then normalized to the 1xCBS_Fw_Sox2P reporter expression at LP-116 kb measured on the same day. Each dot represents a clonal cell line. Bottom: relative *Sox2*::*mCherry* expression of the same clones. Grey vertical lines indicate the position of the two LPs.

In LP-116 kb, which is ∼6 kb upstream of the endogenous *Sox2::mCherry* gene, all reporters showed low reporter expression **(Fig. S3C)**. In this position, reporters with forward CBSs (1xCBS_Fw_Sox2P and 3xCBS_Fw_Sox2P) showed ∼1.8-fold higher expression than the promoter-only reporter (Sox2P), whereas the 3xCBS_Rv_Sox2P reporter was expressed at 70% of the control level (**Fig. 3B**). As expected, reporter constructs integrated in LP-39 kb showed increased expression to various degrees **(Fig. S3C)**. Expression of the Sox2P increased 3.8-fold compared to the less permissive LP-116 kb location. Two clones containing 1xCBS_Fw_Sox2P reached, on average, ∼10-fold higher reporter expression (**Fig. 3B, LP-39 kb**). Clones containing the 3xCBS_Fw constructs showed an average increase of 13.9-fold, with a maximum of ∼25-fold, compared to the Sox2P reporter in LP-116 kb. Inverting the 3xCBS motifs in front of the reporter (3xCSB_Rv_Sox2P) led to a similar increase in reporter expression (∼10-fold) but to a lower maximum activation (14.6-fold) (**Fig. 3B, LP-39 kb**). Overall, these results agree with the hopping-based expression scores at these two locations **(Fig. S3G)**.

We noticed substantial clonal variation in the activity of reporters carrying forward-oriented CBSs inserted in LP-39 kb (**Fig. 3B**). Genotyping the inserts by PCR and Sanger sequencing **(Fig. S3D & S3E)** revealed no genetic alterations. Repeated measuring of these clones over 11 days of culturing showed stable expression of each clone **(Fig. S3F)**. At present we do not have an explanation for this variation between clones **(Fig. 2G).**

### CBS-containing reporters mildly inhibit endogenous Sox2 expression

Next, we assessed the effect of CBS-containing reporters on the expression of the endogenous *Sox2::mCherry* gene. First, we measured *Sox2::mCherry* levels of the same clonal cell lines as shown in **Fig. 3B**. In LP-116 kb, none of the added CBSs impacted *Sox2::mCherry* levels in comparison to Sox2P **(Fig. 3C)**. As we reported previously (*23*), insertion of SoxP in LP-39 kb also did not affect *Sox2::mCherry* levels, despite its location between the endogenous gene and the SCR. Addition of CBSs in this position reduced *Sox2::mCherry* levels by on average 15, 35 and 17% for 1xCBS_Fw_Sox2P, 3xCBS_Fw_Sox2P and 3xCBS_Rv_Sox2P, respectively **(Fig. 3C)**. To further test positional effects, we analysed 1xCBS_Fw_Sox2P insertions at five additional genomic locations. Despite differences in reporter activity, all constructs caused a similar reduction in *Sox2::mCherry* (∼27% on average) relative to the LP-116 kb position **(Fig. 3D)**. These modest reductions are consistent with previously reported CBS insertions (without reporter) in the *Sox2*-SCR interval (*17, 18*). Furthermore, our data suggest that reporter expression levels are not deterministic for the effect on *Sox2::mCherry* levels, because clones with the same insert but different reporter expression display the same effect on *Sox2::mCherry* (**Fig. 3C, 3D**), even when measured over multiple days **(Fig. S3F)**.

### Proximal CBS-promoter arrangement occurs frequently throughout the genome

The results presented above indicate that forward-oriented, but not reverse-oriented, CBSs just upstream of a promoter transcribing in the same orientation can boost activity of the promoter and mildly confine the activation realm of the enhancer. We wondered whether this joint CBS-promoter configuration is common in the genome. Therefore, for each annotated TSS in the mouse genome, we classified the orientation of all nearby predicted CBS motifs into three groups (1) all CBS motifs in the same orientation as the TSSs; (2) all CBS motifs in the opposite orientation as the TSSs; and (3) mix of CBS orientations relative to TSS orientation **(Fig. 4A)**. We then quantified the fractions of these configurations for different genomic window sizes upstream or downstream of the TSSs. As expected, the majority of TSSs does not have a CBS within smaller window sizes of the TSSs (94% upstream and 95% downstream at 2,500 bp) but the number of CBS increases with increasing window size **(Fig. S4A & S4B)**. Nevertheless, when analysing the TSSs with at least one CBS within the set window size, a clear bias emerges upstream of the TSS: same orientation CBSs occur more frequently **(Fig. 4B)**. This bias is most visible within smaller window sizes up to 2,500 bp **(Fig. 4C)**. The distribution of CBS motifs in regards to the orientation and location of the TSSs revealed an enrichment of CBSs upstream of TSSs, in the range of about –750 bp to +200 bp, and in the same orientation **(Fig. 4D)**. Overall, this result suggest that the genome evolved in a way that favours having CBS upstream in the same orientation as transcription. This is in line with our observed CBS-reporter bias in the *Sox2* locus.

**Fig 4.**
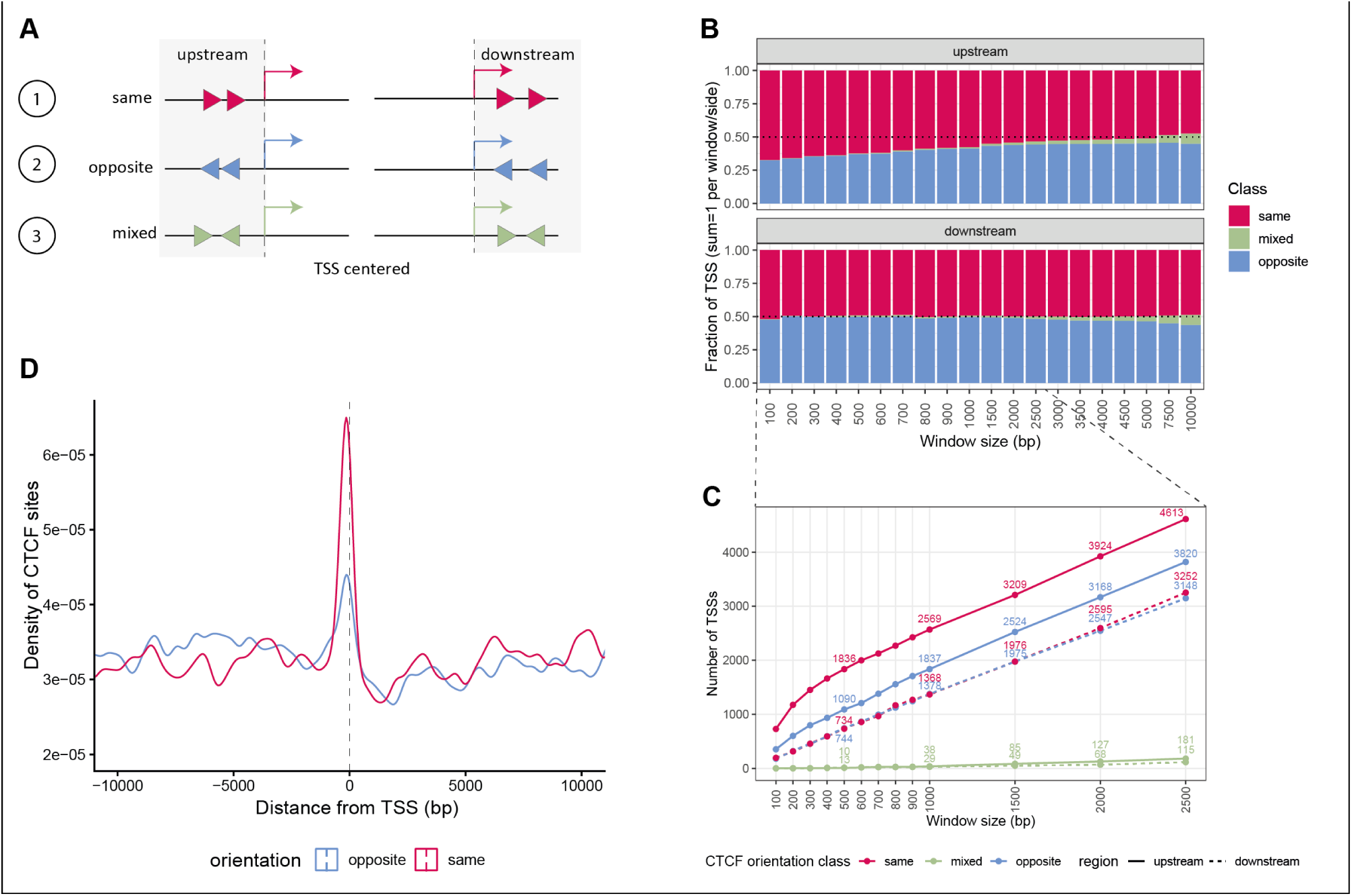
Genome wide orientation and location biases of CBSs in regards to TSSs. **(A**) Cartoon model for classification of CBS-TSS pairs. CBS-TSS pairs can be classified into three categories: (1) all CBSs are in the same orientation as the TSSs (2) all CBSs are in the opposite orientation as the TSSs (3) mix of CBS orientations, either upstream or downstream of the TSSs. **(B)** Fraction of TSSs in different window sizes (100–10,000) that are in the same, opposite or mixed orientation as the CBSs. Analysis was done for CBSs that are upstream or downstream of TSS. Dataset was filtered for having at least one CBS present per window. **(C)** Absolute number of differently classified TSSs for different window sizes. **(D)** Density distribution of the distance (bp) of all putative CBSs to TSSs in a window of +/− 10 kb. Density was calculated with a bandwidth of 250 bp. Colors indicate the orientation of CBS-TSS pairs.

### Location and orientation specific insulation of *Sox2::mCherry* by CBSs

In a second series of “hopping” experiments, we studied how integrated CBSs – individually, not linked to a promoter – might affect the functional interaction between SCR and the endogenous *Sox2* gene. Previous studies have shown that insertion of one or more CBSs between *Sox2* and SCR mildly reduces *Sox2* expression (*17, 18*). However, these studies were limited to a handful of insertion positions; it has remained unclear how the impact of a CBS might depend on its location throughout the locus. Here, we used our SB transposition approach to generate high-resolution maps of the ‘insulation landscape’ of CBSs across the *Sox2* locus.

We designed several constructs for these hopping experiments. One construct simply contained a single reverse CTCF motif (1xCBS), flanked on either side by 25 bp of endogenous sequence **(Fig. 5A)**. This CBS is derived from the Acvr2a locus and has been shown to bind CTCF and cohesin (based on ChIP-seq data, data not shown). The CBS was flanked by loxP sites in either the same (1xCBS_loxP(Fw)) or opposite (1xCBS_loxP(Rv)) orientation, enabling generation of matched ΔCBS control cell lines by Cre-mediated excision **(Fig. 5A)** or Cre-mediated inversions (not used here) of the motif. For this study, we generated a ΔCBS control clone by Cre-mediated deletion of the 1xCBS motif from the 1xCBS_loxP(Fw) cell line before hopping. To maximize insulation, we additionally used the synthetic 3xCBS sequence previously used in the 3xCBS_Sox2P reporters **(Fig 1B & 5A)**. Together, these constructs allowed for a quantitative test of dose– and position dependent insulation.

**Fig 5.**
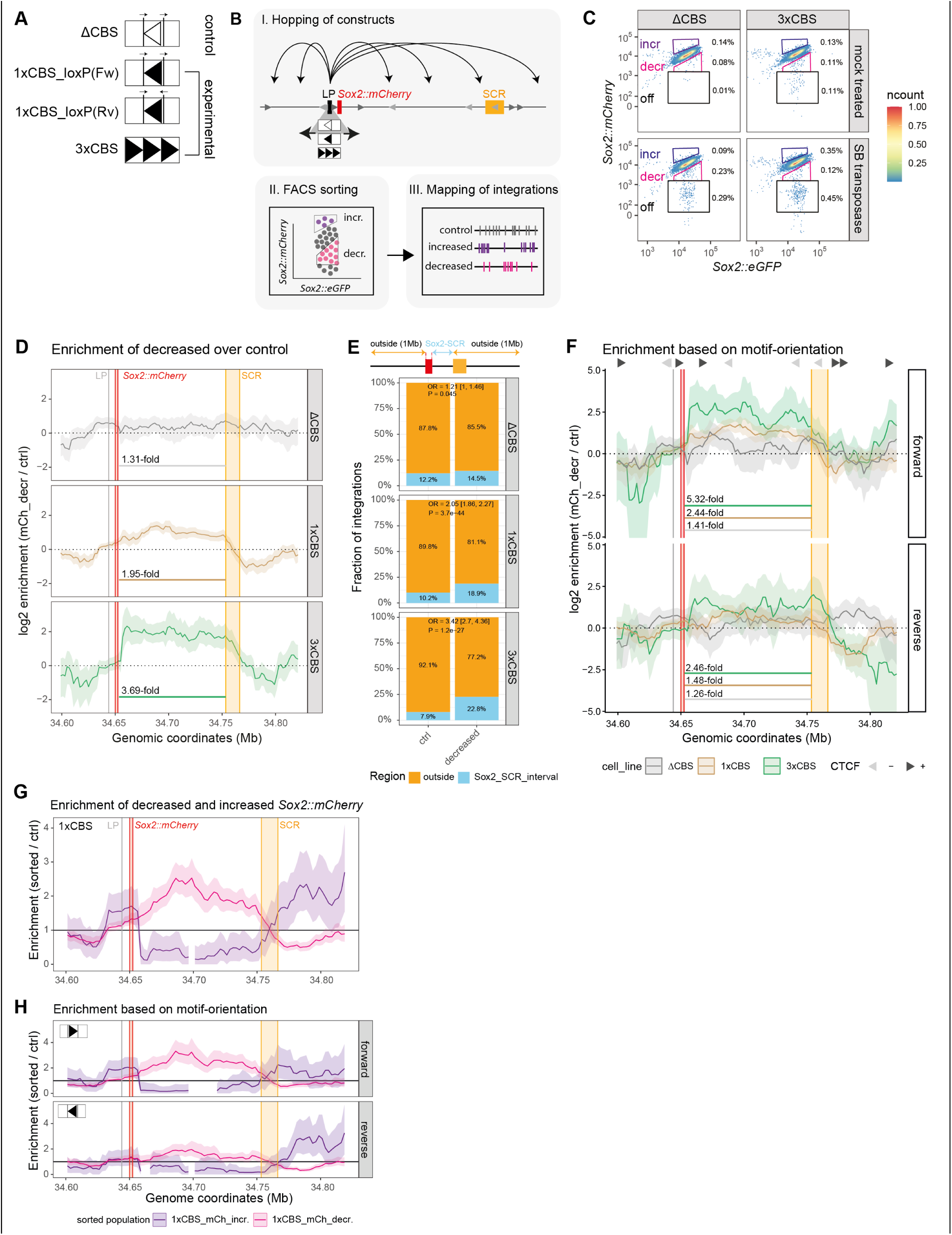
Highly detailed insulation/activation landscapes of CBS in the *Sox2* locus. (**A**) Schematic representation of CBS transposon constructs. Control: ΔCBS construct; Experimental: 1xCBS_loxP(Fw) & 1xCBS_loxP(Rv) constructs. Both differ only in the orientation of the second loxP site; 3xCBS construct contains the same 3xCBS sequence as reported earlier for the 3xCBS_Fw_Sox2P reporter. **(B)** Cartoon depicting the experimental workflow: (1) Initiate hopping of construct from LP-116 kb (2) FACS sort cells for either decreased *Sox2*::mCherry ‘mCh_decr’ or increased *Sox2*::mCherry ‘mCh_incr’ expression (3) map integrations of sorted populations and calculate enrichment (sorted/control) **(C)** Expression levels of *Sox2*::mCherry and *Sox2*::eGFP (control) in mock-transfected (PiggyBac) cells and in cells transfected with SB transposase, measured by FACS. Boxes mark sorting gates used to obtain cell populations ‘mCh_decr’ and ‘mCh_incr’. 200k unsorted control (hopped) cells were collected and mapped to establish normal distribution of integrations. Percentage of cells per gate is indicated. Representative result of multiple replicates. **(D)** Enrichment of integrations sorted for reduction in *Sox2*::mCherry (‘mCh_decr’) over the normal distribution of integrations (unsorted control) per construct. Enrichment was smoothened using a 20kb window, shifted by 2kb steps and only plotted if at least 3 control integrations per window are present. Shaded region indicates 95% CI. **(E)** Fraction of integrations outside or within *Sox2*-SCR interval between control or ‘mCh_decr’ sorted population stratified by construct. For statistical analysis a Fisher exact test was performed and the p-value and odds ratio (OR) are reported. **(F)** Motif-orientation specific enrichment of ‘mCh_decr’ sorted population over control for different constructs. Grey triangles indicate position and orientation of endogenous CBS. Motif-orientation specific enrichment was smoothened using a 20kb window, shifted by 2kb steps and only plotted if at least 3 control integrations per window are present. **(G)** Enrichment of ‘mCh_incr’ and ‘mCh_decr’ sorted integrations over control population. Enrichment was smoothened using a 50kb window, shifted by 10kb steps and only plotted if at least 3 integrations per window are present. Shaded region indicates 95% CI **(H)** Motif-orientation specific enrichment of ‘mCh_decr’ and ‘mCh_incr’ over control.

### High throughput relocation of CBSs

To assess the effect of the different CBS constructs across the *Sox2* locus, we integrated and mobilized them from LP-116 kb **(Fig. 5B)**. We then FACS sorted cells for either decreased (‘mCh_decr’) or increased (‘mCh_incr’) *Sox2::mCherry* expression **(Fig. 5B)**. Control distributions were established for each construct by mapping the integrations of unsorted mobilized pools. Relative to mock-treated controls, hopped pools exhibited broader *Sox2::mCherry* distributions **(Fig. 5C, S4C)**, consistent with position-dependent effects. A small fraction of cells fully lost *Sox2::mCherry* (off gate; **Fig. 5C & S4C**). In initial experiments we found that this is predominantly the result of deletions of the entire gene, which occur due to rare events of aberrant SB hopping (*23*). Because we were interested in the effect of CBSs on *Sox2* insulation rather than deletions, we excluded cells with such complete loss of *Sox2::mCherry* expression from further analysis.

To address the positional effect of CBSs on the reduction of *Sox2::mCherry* (‘mCh_decr’), we performed multiple independent replicates for ΔCBS (3x), 1xCBS_loxP(Fw) (3x), 1xCBS_loxP(Rv) (2x) and 3xCBS (2x). The integration patterns between replicates of the cell lines were reproducible **(Fig. S4D-S4G)**; therefore, we combined the replicates for further downstream analysis. For 1xCBS, we initially generated two clonal cell lines containing the 1xCBS reverse motif. Those clones only differ in the arrangement of loxP sites surrounding the CBS. Based on a strong correlation of baseline hopping distribution (Pearson r = 0.8, 5 kb bins; **Fig. S4H**) between both clones, we combined them for further analysis. In total, we mapped 1,182 (ΔCBS), 4,685 (1xCBS) and 1,292 (3xCBS) independent integrations +/− 1Mb from LP-116 kb (**Fig. S5A**) from *Sox2::mCherry* reduced cells, in each case with at least 446 integrations distributed across the 240 kb region around the *Sox2*-SCR interval (**Fig. S5B**).

To visualize position-dependent effects, we calculated the enrichment of cells with decreased Sox2::mCherry expression (‘mCh_decr’) over the unsorted control integrations, using a running window across the locus **(Fig. 5D)**. As expected, the sorted ΔCBS control produced a near-flat enrichment profile across the locus **(Fig. 5D)**, with only a marginal enrichment of integrations (odds ratio OR = 1.21, p = 0.045) in the *Sox2*-SCR interval compared to unsorted hopping events **(Fig. 5E)**. This indicates that the SB transposon without the CBS has rarely any impact on *Sox2::mCherry* expression throughout the locus and the sorted cells were the result of stochastic fluctuations in mCherry expression. In contrast, the resulting integration landscapes of CBS integrations (1xCBS & 3xCBS) show a clear enrichment throughout the interval, indicating that the insulation effect is not dependent on the exact genomic location within this interval. 1xCBS integrations sorted for *Sox2::mCherry* reduction were ∼2-fold enriched compared to the unsorted control within the *Sox2*–SCR interval, flanked by sharp transitions near the *Sox2* gene and downstream of the SCR **(Fig. 5D)**. Consistently, the fraction of integrations within the *Sox2*–SCR interval increased from 10% (control) to 19% (‘mCh_decr’) (p = 3.7×10⁻^44^, OR = 2.05; **Fig. 5E)**. Analysis of the 3xCBS construct showed an even more pronounced confinement, with ∼3.7-fold enrichment across the *Sox2*–SCR interval **(Fig. 5D)** and an increase from 8% (control) to 23% (‘mCh_decr’) (p = 1.2×10⁻^27^, OR = 3.42; **Fig. 5E**). Together, these data illustrate that CBSs decrease *Sox2::mCherry* expression in a dose– and location-dependent manner, with maximal effects between *Sox2* and the SCR.

To test whether these CBS effects depend on motif orientation, we stratified integrations by orientation **(Fig. S5B)** and recalculated the enrichments. The ΔCBS control showed almost no orientation bias within the *Sox2*–SCR interval in cells sorted for mCherry reduction compared to the non-sorted control population **(Fig. 5F)**. Furthermore, the fractions of integrations within this interval are almost equal (forward 14.3% vs reverse 14.6%; **Fig. S5C**). In contrast, 1xCBS showed a mean 2.4-fold increase in forward motif integrations within the *Sox2*-SCR interval while the reverse motif integrations were almost equal (1.48-fold) to the ΔCBS control cell line (1.26-fold). For 3xCBS, both motif orientations were enriched compared to the ΔCBS control, whereas forward motif integrations have a 5.3-fold higher enrichment throughout the *Sox2*-SCR interval compared to 2.5 for reverse motif integrations **(Fig. 5F)**.

Next, we analysed the rare integration patterns of the 1xCBS and 3xCBS constructs in cells that displayed increased (‘mCh_incr’) *Sox2*::mCherry expression **(Fig. 5C)**. We performed two replicates for 1xCBS_loxP(Fw), one replicate for 1xCBS_loxP(Rv) and one replicate for 3xCBS. Integrations from 1xCBS clones were again combined for further analysis. This resulted in 432 and 75 mapped ‘mCh_incr’ integrations within +/− 1Mb from LP-116 kb for 1xCBS and 3xCBS, respectively. These integrations almost exclusively mapped outside the *Sox2*-SCR interval **(Fig. S5A & S5B)**. When we compare the enrichment landscape from the 1xCBS cell line sorted for increased or reduced *Sox2*::mCherry (‘mCh_incr’ & ‘mCh_decr’), a clear difference in enrichment patterns emerges. CBS integrations leading to increased *Sox2*::mCherry expression are almost absent within the *Sox2*-SCR interval and become enriched downstream of the SCR **(Fig. 5G)**. The same can be seen for the 3xCBS cell line **(Fig. S5D)**. Stratifying the integrations by CBS motif-orientation indicates that both motif-orientations are enriched downstream of the SCR, with the reverse motif orientation having a stronger enrichment in both cell lines **(Fig 5H, Fig S5E)**. Nevertheless, motif-orientation specific interpretations should be taken with caution because of the low number of integrations leading to large confidence intervals.

In summary, relocation of CBS constructs to thousands of locations revealed a dosage-dependent inhibitory effect of CBS throughout the *Sox2*-SCR interval. This effect is more pronounced when the CBS is oriented towards the SCR. Upregulation of the gene seems to be enriched for reverse oriented CBS integrations downstream the SCR. While *Sox2* regulation has been reported to be relatively loop-extrusion independent in the wild-type locus configuration (*27*), our hopping approach detects subtle, orientation-linked expression shifts, consistent with a constrained and asymmetric insulation landscape across the locus.

### Hopping distributions suggest that CBS-containing reporters alter 3D folding

Upon closer inspection of the integration patterns for the different inserts in the two series of hopping experiments **(Fig. 6A)**, we noticed systematic differences in their genomic distributions across the locus. Inserts carrying forward-oriented CBS motifs in LP-116 kb (1xCBS_Fw_Sox2P, 3xCBS_Fw_Sox2P and 3xCBS_Fw) displayed a clear bias toward downstream integrations compared to non-CBS controls (Sox2P, ΔCBS) **(Fig. 6B & 6C)**. In contrast, transposons carrying reverse-oriented CBS motifs in LP-116 kb (3xCBS_Rv_Sox2P, 1xCBS_Rv) showed a pronounced bias towards integrations upstream of the LP **(Fig. 6B & 6C)**. Quantitative analysis confirmed this: among the unsorted integrations, 1xCBS_Fw_Sox2P, 3xCBS_Fw_Sox2P and 3xCBS_Fw showed higher odds of inserting downstream than upstream of the LP (odds ratios OR = 2.0, 2.7 and 2.5, respectively), whereas reverse-oriented constructs (3xCBS_Rv_Sox2P and 1xCBS_Rv) displayed lower odds than their respective controls (OR = 0.382 and 0.767, respectively; **Fig. 6D**).

**Fig 6.**
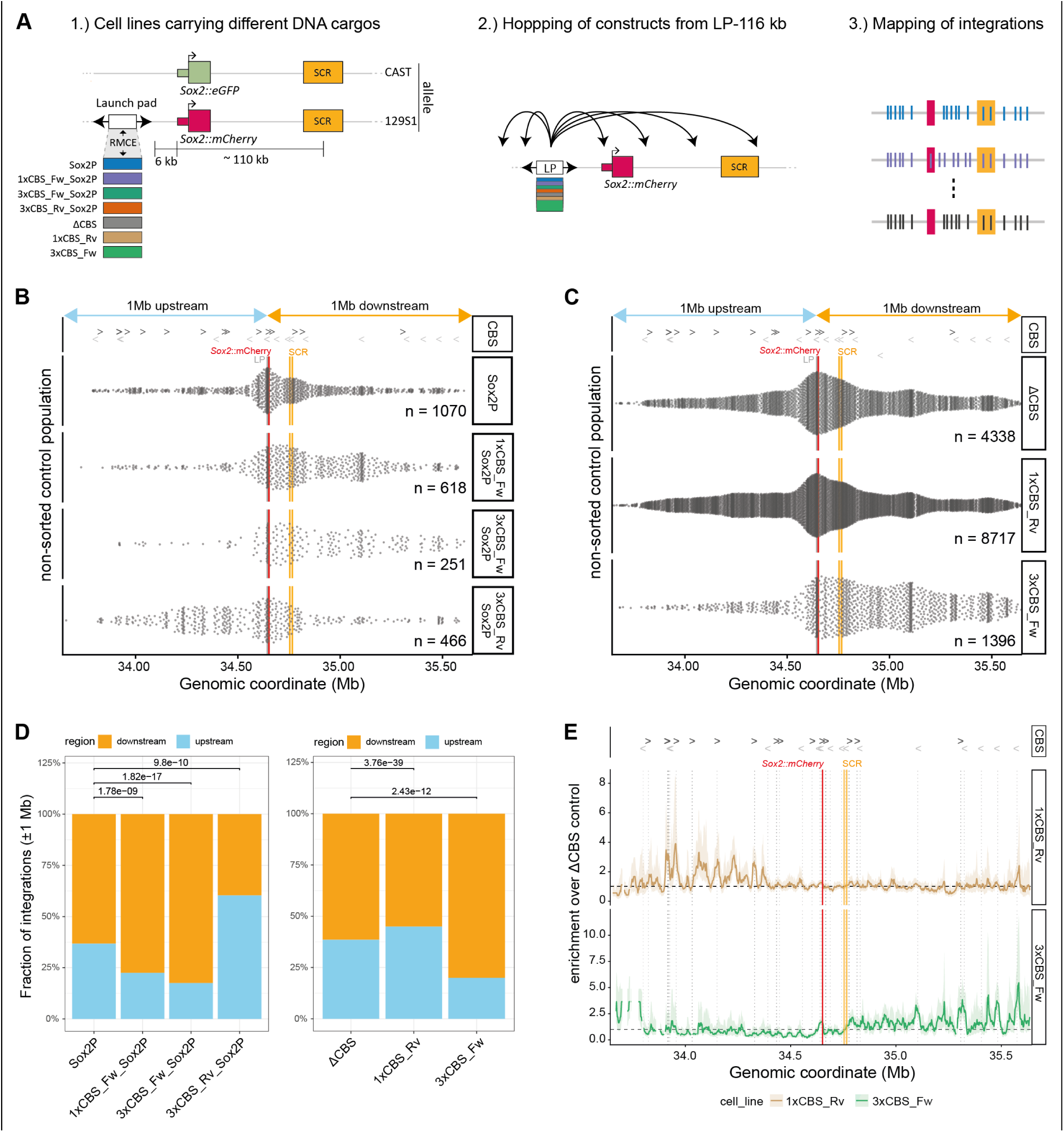
Transposon ‘cargos’ can influence hopping behavior. (**A**) Schematic representation of the experimental workflow. (1) Integration of different transposon cargos in LP-116 kb. (2) Hopping of constructs from LP-116 kb. (3) Mapping the unsorted control integrations from each construct. **(B)** Distribution of unsorted control integrations (+/− 1Mb from LP) containing different reporter constructs hopped from LP-116 kb. Each dot represents a mapped integration. Number of integrations plotted is indicated by n. Location and orientation of endogenous CBS (CBS) are indicated with arrows. **(C)** same as (B), except different transposon constructs are displayed. **(D)** Fraction of control integrations that are upstream (1Mb upstream of LP-116 kb) or downstream (1Mb downstream of LP-116 kb) of LP-116 kb per transposon construct. Fisher Exact test was used to determine significant differences between Sox2P (control) and 1xCBS_Fw_Sox2P, 3xCBS_Fw_Sox2P and 3xCBS_Rv_Sox2P, respectively. The same was done for the CBS only constructs by comparing ΔCBS (control) to 1xCBS_Rv and 3xCBS_Fw. **(E)** Enrichment of CBS containing integrations (1xCBS_Rv, 3xCBS_Fw) over the ΔCBS control cell line was calculated using a smoothened 20kb window, shifted by 2kb steps and only plotted if a least three integrations per window were present for each cell line. Shaded area indicates 95% CI.

Moreover, all CBS-containing inserts showed a pronounced enrichment of integrations near endogenous CBS sites. Calculating the enrichment of integrations of the 1xCBS_Rv and the 3xCBS_Fw constructs over the ΔCBS control revealed clear peaks of enrichment near endogenous CBSs upstream or downstream of the launch pad, respectively **(Fig. 6E)**. The same trend was observed for the CBS-containing Sox2P reporters **(Fig. S5F)**. Together, these results point to substantial “homing” of CBS-containing SB transposons, i.e., an increased probability of transposition into the vicinity of another CBS. Because this homing primarily involves convergent CBSs, we propose that cohesin-mediated loop extrusion (*4, 5*) underlies this process: a loop is likely formed prior to SB excision, increasing the chance that the transposon hops into DNA that is brought into spatial proximity by this loop. Additional evidence that SB hopping is at least partly driven by spatial proximity will be reported elsewhere (manuscript in preparation) and was also noted for LAD versus inter-LAD compartmentalization (*28*) and in the context of polycomb-mediated pairing of distant loci (*29*). Besides enrichment at endogenous CBSs, there seem to be more factors influencing the CBS-containing inserts hopping behaviour, e.g. non– annotated CBSs or other contact facilitating elements, displayed by peaks of integrations not overlapping endogenous CBSs.

We emphasize that, while these hopping biases affect the local *density* of integrations, they do not skew the calculation of the *levels* of the expression scores (see **Methods**); they only affect the local *accuracy*, which generally increases with higher integration densities in the sliding window that is used for the calculation. Moreover, the fraction of reporters that hopped into the *Sox2*-SCR interval – the region with the most pronounced functional differences between the reporters – is almost equal between the different reporter constructs, ranging from 12% (Sox2P) to 15% (1xCBS_Fw_Sox2P) **(Fig. S5G)**, and within the *Sox2*-SCR interval we did not observe strong local enrichments of insertions at CBSs (**Fig. 6E**).

### Deletion of CBS in/near SCR leads to lower reporter expression

In a third series of “hopping” experiments we dissected the impact of two endogenous CBSs on the regulatory landscape of the *Sox2* locus. One CBS, located in the middle of the SCR and oriented towards the *Sox2* promoter, was previously reported to enhance the contact frequency between *Sox2* and SCR (*11*). Although deletion of this CBS did not detectably alter the expression level of *Sox2* (*11*), we hypothesized that this CBS might modify the activation landscape elsewhere in the locus. Another CBS that drew our attention was the CBS located ∼12 kb downstream of the SCR, coinciding with a sharp transition from active to silent Sox2P reporters **(Fig. 2B).** We hypothesized that this CBS might insulate the downstream region from activation by the SCR. Here, we studied the effect of deleting each of these two CBSs on the overall regulatory landscape of the locus by large-scale hopping and mapping of the Sox2P reporter.

We used Cas9 editing to delete each CBS in separate cell lines (ΔCBS-in-SCR and ΔCBS-DS[downstream]-SCR) containing the Sox2P reporter in LP-116 kb upstream of the SCR (**Fig. 7A, S6A-S6B**). Deletion of each CBS independently reduced Sox2P reporter expression by 30-50%, without affecting the expression of the endogenous *Sox2::mCherry* gene (<2%) (**Fig. 7B**). The reduction in reporter expression is a direct effect of deleting the CBS on the same allele as the reporter (129S1), since a clone with a heterozygous deletion in the CAST allele (clone D13) displays reporter expression equal to non-edited cells **(Fig. S6C, clone ΔCBS-in-SCR_D13)**.

**Fig 7.**
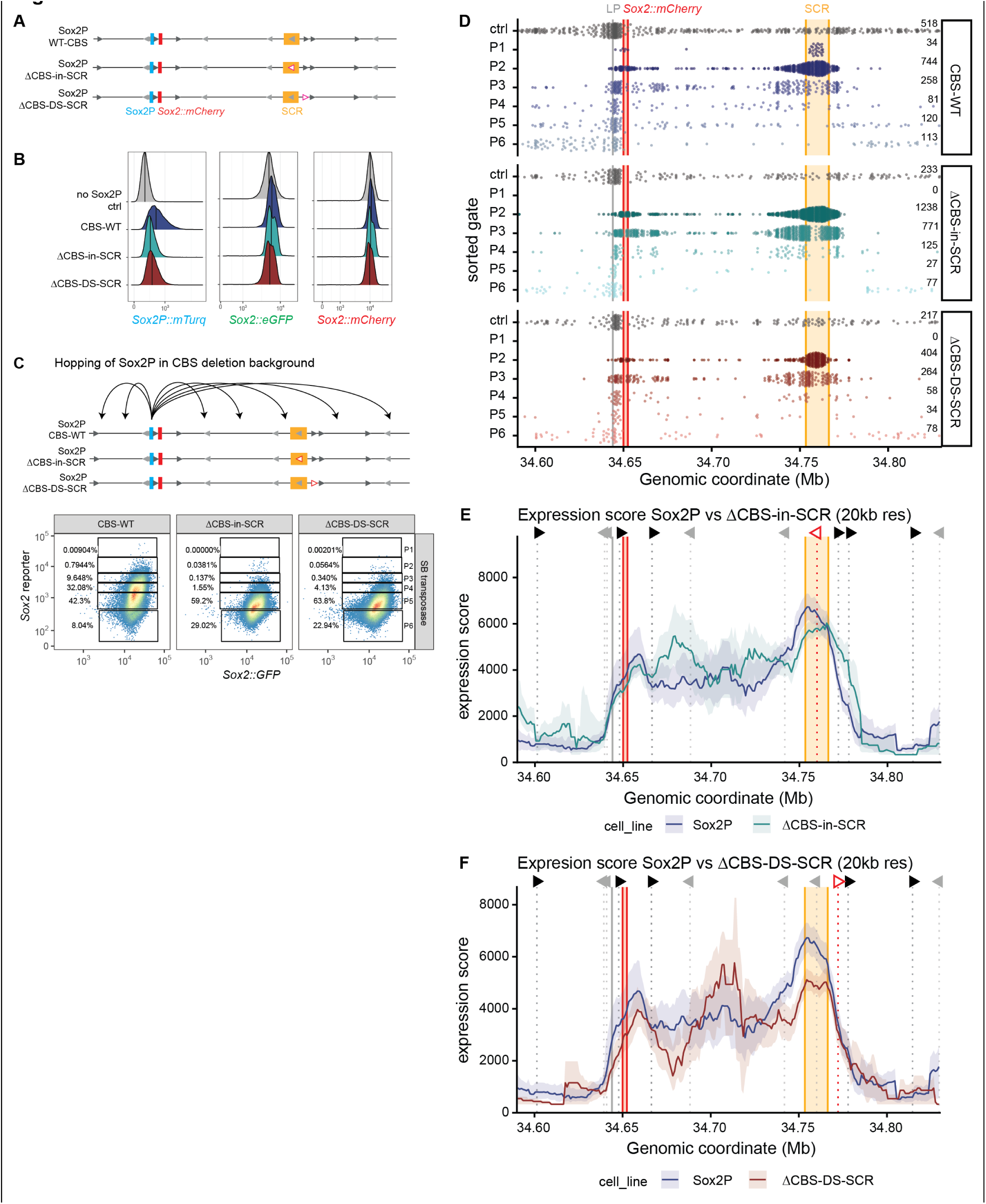
Effect of endogenous CBS on reporter expression and enhancer reach. (**A**) Schematic representation of control (Sox2P_WT-CBS) and edited (Sox2P_ΔCBS-in-SCR & Sox2P_ΔCBS-DS-SCR) cell lines containing the Sox2P reporter in LP-116 kb. Blue rectangle: Sox2P reporter; red: *Sox2*::mCherry; green: *Sox2*::eGFP; orange: SCR; position and orientation of grey triangles indicate endogenous CBS; pink triangles indicate CRISPR/Cas9 deleted CBSs. **(B)** Endogenous *Sox2* levels (*Sox2*::eGFP & *Sox2*::mCherry) and Sox2P reporter expression (*Sox2*::mTurq) of control (non-fluorescent control and CBS-WT control) and CBS deletion cell lines (Sox2P_ΔCBS-in-SCR & Sox2P_ΔCBS-DS-SCR) containing the Sox2P reporter in LP-116 kb measured by FACS. **(C)** Top: Schematic of hopping the Sox2P reporter; Bottom: Sox2P reporter and *Sox2*::eGFP FACS measurements of SB transfected cells. Reporter expression is divided into six gates (P1-P6) and cell pools were sorted for each gate. Percentage indicates the frequency of cells occurring per population. **(D)** Distribution of mapped integrations per cell line (CBS-WT; Sox2P_ΔCBS-in-SCR, Sox2P_ΔCBS-DS-SCR) and sorted population (P1-P6). Each dot represents a mapped integration. The number of integrations plotted per population is indicated by n. Ctrl integrations refer to the mapping of a large pool of unsorted (hopped) cells. Grey vertical line: position of LP-116 kb; red vertical square: position of *Sox2::mCherry*; orange vertical square: position of the SCR. **(E)** Expression score calculate of Sox2P integrations mobilized from LP-116 kb in CBS-WT (ctrl) and ΔCBS-in-SCR in (D), smoothened using a 20kb window shifted by 1kb steps and only plotted if at least three integrations per window are present. Shaded region indicates 95% CI. Grey vertical dotted lines and triangles indicate the position and orientation of CBSs. **(F)** Same as (E), except the comparison with cell line ΔCBS-DS-SCR is displayed.

We then induced Sox2P reporter hopping, monitored reporter fluorescence by flow cytometry and sorted cell pools from six gates as previously described **(Fig. 7C)**. Few cells reached the highest reporter expression level (P1) in each cell line, but across replicates a mean of 0.007 % (0.003 – 0.009 %) of Sox2P CBS-wildtype (Sox2P_CBS-WT) cells were in this gate, while in both CBS deletion cell lines we observed on average (across replicates) one out of 100k cells (0.001 %) in P1 (**Fig. 7C, FACS plots**). This suggests that deletion of individual CBS can prevent maximum reporter activation in certain locations. After mapping, the integration patterns showed a good correlation between biological replicates of the respective cell lines; hence we combined the data for further downstream analysis (**Fig. S7A & S7B**). The Sox2P CBS-WT control data is the same data as illustrated in **Fig. 2C, 2D & 2E.** Within the 240 kb region around *Sox2*-SCR, we obtained more than thousand integrations for each cell line **(Fig. 7D)**.

The mapped integrations revealed location-dependent reporter expression **(Fig. 7D)**. All three cell lines showed highest activation of the reporter around the endogenous *Sox2::mCherry* gene or within the SCR. As indicated earlier, no integrations corresponding to the highest reporter expression (P1) were obtained from the CBS deletion cell lines (ΔCBS-in-SCR, ΔCBS-DS-SCR). Intermediate reporter expression (P3, P4) integrations are depleted in the core SCR region and redistribute downstream of *Sox2* and upstream of the SCR, whereas silent reporters were positioned outside the TAD **(Fig. 7D)**. The reporter expression score tracks **(Fig. 7E & 7F)** appeared globally similar between the CBS deletion cell lines and CBS-WT, with intermediate reporter activity throughout the *Sox2*-SCR interval and sharp transitions just upstream of *Sox2* and downstream of the SCR. In agreement with the missing P1 integrations, CBS deletion cell lines showed lower reporter activation at the SCR and the endogenous *Sox2::mCherry* gene **(Fig. 7E & 7F).** Investigating the boundaries more locally, it appears that deletion of the CBS-in-SCR leads to a spread of reporter activation downstream of the SCR until encountering the next endogenous CBS **(Fig. 7E)**. Deletion of the CBS downstream of the SCR did not lead to such spreading of reporter activity but instead further restricts reporter activation at the endogenous gene **(Fig. 7F)**.

Overall, these data indicate that deleting CBSs in the centre or downstream of the SCR can have substantial effects on maximum reporter expression throughout the locus whereas the endogenous *Sox2* gene is unaffected by the loss of either CBS, suggesting that the reporters are more sensitive to structural changes in the locus. The endogenous *Sox2* gene might be insensitive to this perturbations because of its optimal position in the genome, right next to regulatory elements (SRR2) (*27*) and because it contains the 1kb coding sequence, which has a boosting effect on gene expression (*23*). Deletion of CBSs in the SCR or downstream of the SCR indicate that CBSs could act as borders for reporter activation. Assuming that CBSs have a redundant function, the genomic distance between the deleted CBSs and the following one is very small, which makes it difficult to gather enough data to make solid conclusions. Deletion of multiple CBSs at the TAD boundary with subsequent reporter hopping could shed light on the border function of CBSs.

## Discussion

We previously established a transposon-based method to map the regulatory landscape of the endogenous *Sox2* locus by relocating a *Sox2* promoter reporter (Sox2P) to thousands of alternative positions. This approach not only revealed the position-dependent activation landscape in the presence of the endogenous *Sox2* gene but also that the gene (in combination with its upstream CBS) confines the activation realm of the enhancer (*23*). Here, we applied several variations of this technology to test the effect of CBSs on the functional landscape of the *Sox2* locus.

The CBS–promoter constructs that we tested displayed strong orientation– and dosage-dependent differences in responsiveness to the SCR across the *Sox2*–SCR interval. Reporters in which the CBS orientation was aligned with the promoter orientation exhibited the highest activity when integrated in a configuration pointing towards the SCR **(Fig. 8A)**, but little activity when pointing away from the SCR **(Fig. 8B)**. Importantly, these differences cannot be explained by CBS orientation alone. Inverting only the CBS upstream of the reporter **(Fig. 8E & 8F)** did not recapitulate the effects that we observed when the entire CBS–promoter construct was inverted, indicating that reporter activation depends on the combined orientation of the CBS and the promoter. A recent study found that RNA polymerase II can act as a barrier to loop extrusion upon head-on collision with extruding cohesin (*30*). Random integration of a 2 kb DNA fragment containing both a forward-oriented CBS and a TSS demonstrated that TSSs preferentially fold local chromatin along the direction of transcription, whereas CBSs primarily engage in contacts with convergent oriented CBSs (*31*). Furthermore, promoter-proximal CBSs have been shown to facilitate promoter-enhancer search in three-dimensional space (*8*). Together, these observations suggest that transcription from the Sox2P reporter may synergize with the CBS to impede cohesin extrusion, provided that they are oriented in the same direction.

**Fig 8.**
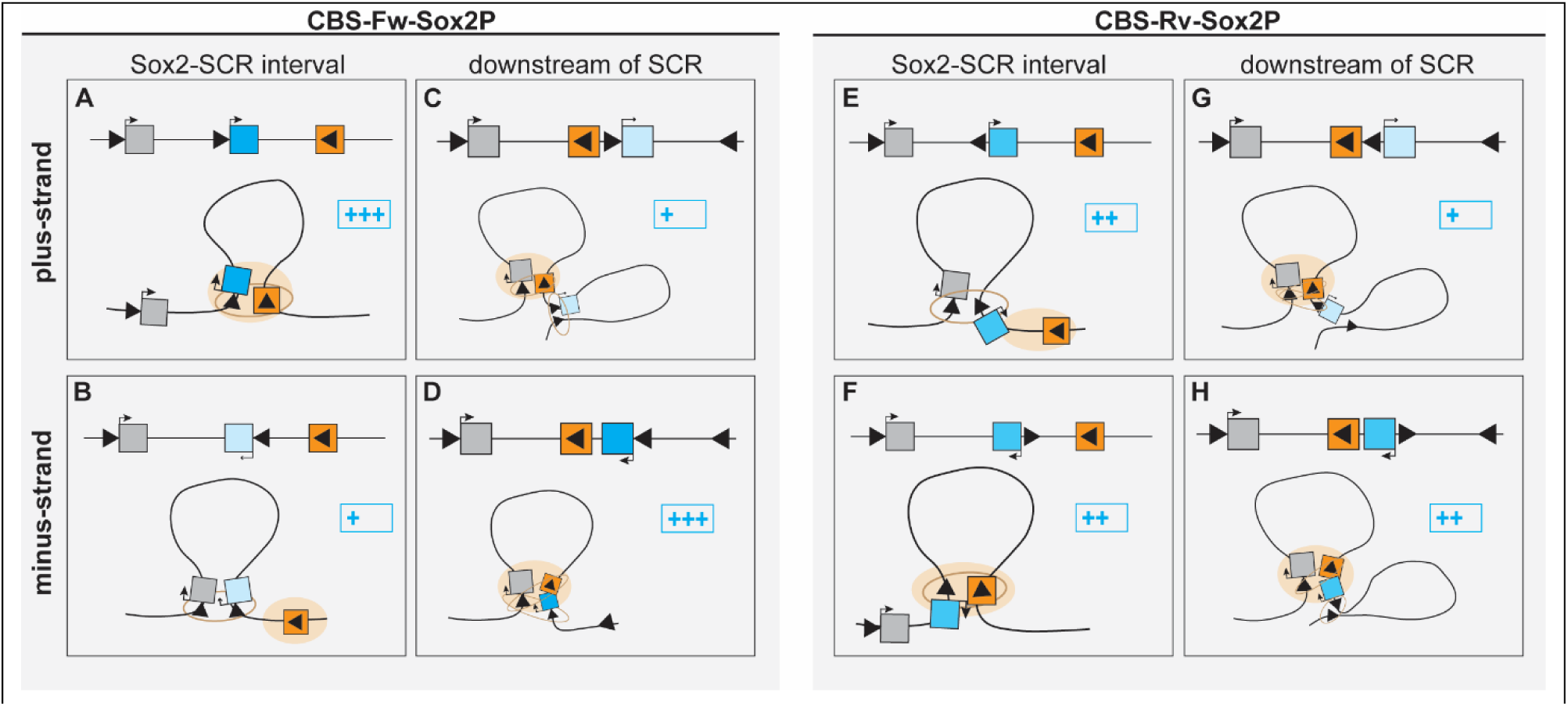
Schematic representation of CBS-Sox2P reporter integrations in context of loop extrusion. *Top row*: CBS-Sox2P reporter integrations in plus-strand; *Bottom row*: CBS-Sox2P reporter integrations in minus-strand. **(A)** Cartoon model of aligned CBS-Fw-Sox2P reporter integrated in between Sox2-SCR interval on the plus or minus **(B)** strand. **(C)** CBS-Fw-Sox2P reporter integrated downstream of SCR on the plus or minus **(D)** strand. **(E-H)** same as A-D, with the CBS-Rv-Sox2P reporter construct instead. Triangles represent position and orientation of CBSs; Blue rectangles represent CBS-Sox2P reporters; Orange rectangles represent SCR; Grey rectangle represents endogenous Sox2 gene and orange circle represent the activation influence conferred by the SCR. Blue plus represent indication for measured reporter activity.

Besides the strong orientation bias observed within the *Sox2*-SCR interval, we observed a switch in motif orientation bias towards the end and downstream of the SCR. Highest reporter activation was restricted to CBS-Sox2P configurations in which the CBS was not in between the SCR and the Sox2P **(Fig. 8D & 8H)**. Changing the orientation so that the CBS is between the SCR and Sox2P **(Fig. 8C & 8G)** reduced reporter expression, indicating that the reporter-proximal CBS can act as a true insulator of enhancer driven promoter activation.

The enrichment of forward-oriented CBSs upstream of promoters in the mouse genome suggest that this CBS–promoter synergy may play a role in the regulation of thousands of genes. Extrapolation of our results suggests that this configuration primarily boosts promoter activity when the enhancer is located downstream, while it may shield the promoter from activation by upstream enhancers. Thus, this configuration may substantially restrict enhancer – promoter interactions at many genomic loci.

The pronounced orientation-dependent asymmetry observed with our CBS-Sox2P reporter and CBS-only integrations may have multiple explanations. One possibility is that loop extrusion traverses the locus more frequently from the SCR side than from the *Sox2* promoter side. This interpretation is underscored by our CBS-only hopping experiments, which show stronger insulation of the endogenous *Sox2*::mCherry gene in the plus strand than in the minus strand. Such a motif-orientation bias was previously observed with CBS integrations at individual positions in the *Sox2*-SCR interval (*17, 18*) and also in two other loci (*19, 32*). Importantly, integrations throughout the *Sox2*-SCR interval show this effect with a similar magnitude. This consistency suggests that loop extrusion initiates more frequently near the SCR, rather than randomly throughout the interval. In the latter case a gradient in the bias would be expected from the SCR towards the *Sox2* gene, which is not what we observed. This preferred initiation near the SCR contrasts with global analyses and simulations suggesting that cohesin might initiate loop extrusion randomly throughout the genome (*33*). However, it is possible that specific regions such as the *Sox2* locus exhibit preferred initiation near the enhancer.

Our data do not rule out alternative models. One other possibility is that the asymmetry results from differential looping strength between the endogenous CTCF sites at the *Sox2* or SCR boundary with the ectopically inserted CBS’s. The enrichment of forward-oriented CBS-only motifs throughout the *Sox2*-SCR interval could result from a stronger interaction with the CBS boundary at the SCR site than with the *Sox2* promoter side.

Deletion of two individual endogenous CBSs within or downstream of the SCR led to a modest reduction in Sox2P reporter expression across the locus, while leaving endogenous *Sox2* expression largely unaffected. This indicates that these CBSs contribute to the efficiency and spatial confinement of SCR-driven activation but are not dominant determinants of *Sox2* transcription. This is consistent with prior observations that *Sox2* expression is relatively insensitive to local perturbations of CTCF-mediated chromatin architecture (*11, 17, 18*). The reporter was substantially more sensitive than the endogenous gene to CBS deletions. Most likely, other genetic elements such as the coding sequence (*23*) or the proximal SRR2 element (*27*) add robustness to the expression of the endogenous *Sox2*. In addition, deletion of the CBS located within the SCR caused a slight expansion of the realm of activation, which terminated at the next endogenous CBS. This indicates that SCR confinement is distributed across multiple architectural elements rather than enforced by a single dominant boundary.

Here, we demonstrate the utility of our hopping approach to study various roles of CBSs, and to obtain indications of the direction of loop extrusion in a model locus. Hopping experiments after simultaneous disruption of multiple CBSs may reveal possible redundancy of these elements. Additional experiments in cells with relocated CBS or hopped CBS-SoxP reporters, such as direct mapping of the 3D contacts, will aid in the deduction of the most likely mechanistic explanation of the results. A disadvantage of the *Sox2* gene is that its expression is only partially dependent on cohesin (*34–36*), which may account for relatively subtle effects of CBSs deletions and insertions on *Sox2* expression. Deletion of the SRR2 element, which was recently found to render the *Sox2* gene partially independent of cohesin (*27*), may expose more pronounced patterns. Obviously, many other genomic loci and regulatory elements will be of interest to explore with this methodology.

**Fig S1.**
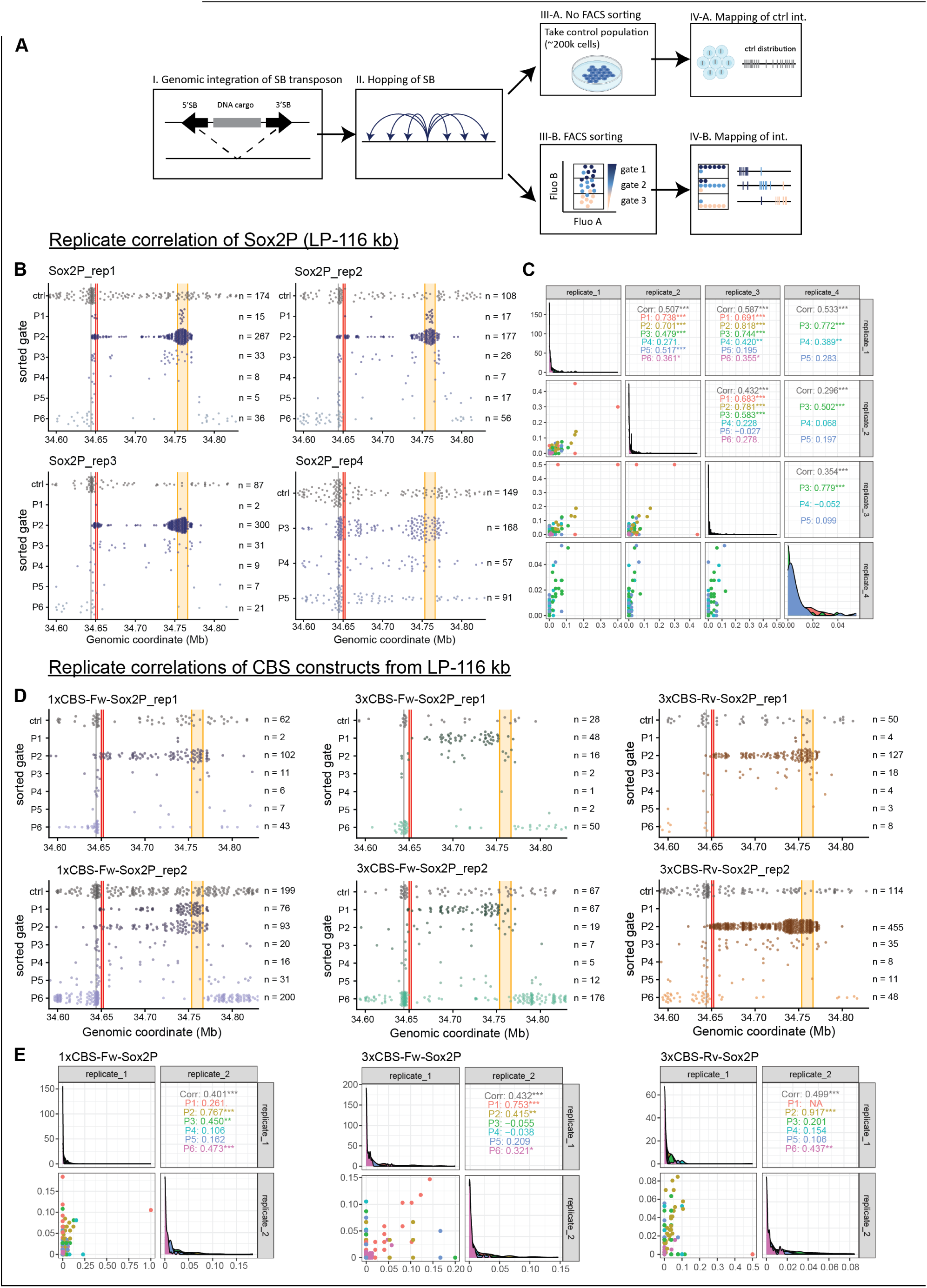
Reproducibility of construct specific hopping from LP-116 kb. **(A)** Schematic representation of hopping workflow. (I) SB transposon gets integrated via CRISPR/Cas9 into location of interest; (II) After loading transposon with cargo, hopping is initiated via SB transposase; (III-A) a large pool of transfected, unsorted cells gets harvested and integrations get mapped; (III-B) transfected cells get sorted into pools of cells based on reporter fluorescence by FACS and integrations for each sorted pool get mapped. **(B)** Distribution of Sox2P integrations from LP-116 kb of four biological replicates across the native *Sox2* locus. For replicate four, 50k cell pools of P3, P4 and P5 populations were sorted and mapped. Each dot represents a mapped integration. Number of integrations plotted is represented by n. **(C)** Reproducibility of reporter integrations across the *Sox2* locus between biological replicates. Scatterplots show the fraction of population integration per 5kb bin (on the interval in Fig S1B) colored by sorted population, across four replicates from LP-116 kb. Diagonal panels show distributions per replicate; upper panels show Spearman correlation coefficients. **(D)** Same as (B) for hopping 1xCBS_Fw_Sox2P, 3xCBS_Fw_Sox2P and 3xCBS_Rv_Sox2P constructs from LP-116 kb. **(E)** As in (C), for 1xCBS_Fw_Sox2P, 3xCBS_Fw_Sox2P and 3xCBS_Rv_Sox2P constructs; two biological replicates each.

**Fig S2.**
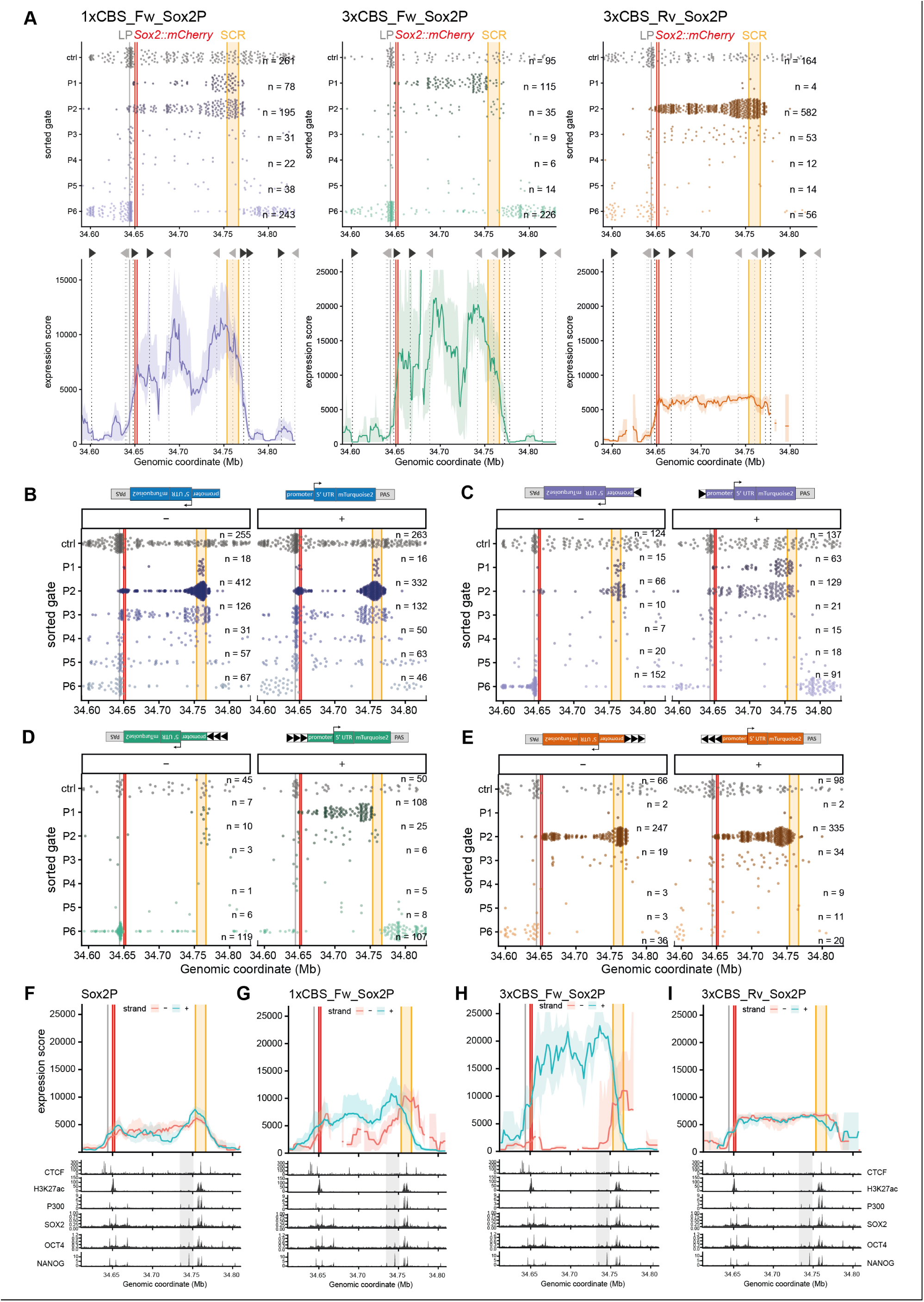
Strand specific effect of CBS reporters on reporter expression. **(A)** Top: Distribution of mapped integrations per sorted population (P1-P6) hopped from LP-116 kb at the *Sox2* locus. Colors indicate the different reporter constructs. Each dot represents a mapped integration. Number of integrations plotted is represented by n. Bottom: Reporter expression scores of the raw integration data displayed at the top are smoothened using a 10kb window shifted in 1kb steps. Shaded region indicates 95% CI. Expression score is only plotted when window contains 3 or more SB integrations. Grey triangles illustrate position and orientation of endogenous CBS. **(B-E)** Distribution of mapped integrations by strand. Cartoons above the scatterplots indicates the orientation of the constructs per strand. **(F-I)** Top: Same stranded reporter expression scores as shown in Fig 2E-H, but colored by strand. Bottom: H3K27ac ChIP-seq P-value signal (*37*), CTCF (*23*), P300 (*38*), SOX2 (*39*), OCT4 (*39*) and NANOG (*39*) ChIP-seq coverage.

**Fig S3.**
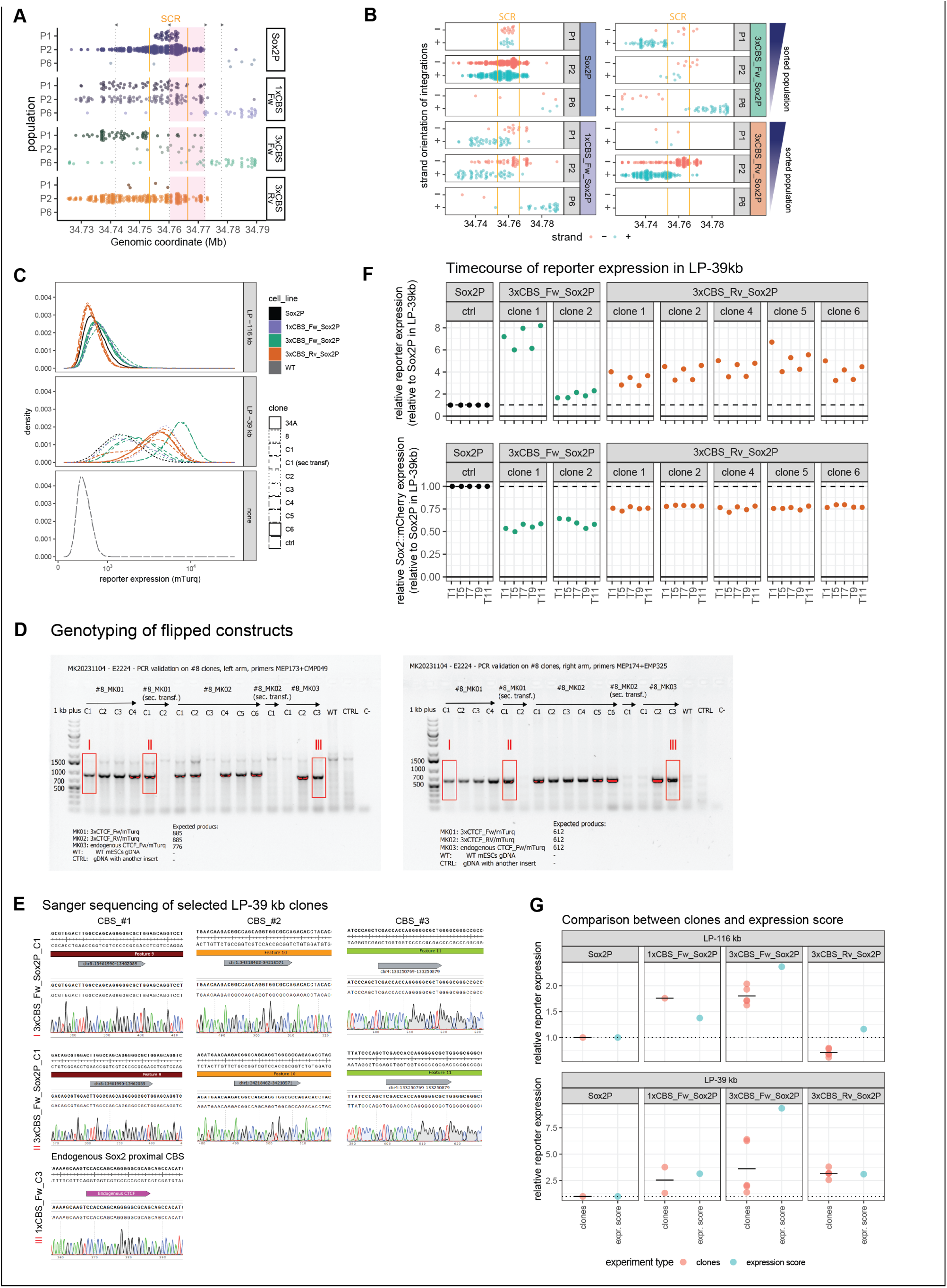
Impact of CBS containing reporters in pools and clones. (**A**) Distribution of mapped integrations from active (P1 & P2) and inactive (P6) reporter populations around the SCR for the four different reporter constructs. Each dot represents a mapped integration. Same data as displayed in Fig 2C and S2A top, zoomed into the SCR region. **(B)** Strand specific distribution of mapped integrations from active (P1 & P2) and inactive (P6) reporter populations around the SCR for the four different reporter constructs. Same data as displayed in Fig S2B-S2E. **(C)** Reporter expression (mTurq) levels of clones containing the four different reporter constructs (Sox2P, 1xCBS_Fw_Sox2P, 3xCBS_Fw_Sox2P, 3xCBS_Rv_Sox2P) integrated in two different launch pads; LP-116 kb and LP-39 kb, measured by FACS. Colors indicate the different reporter constructs; darkblue: Sox2P, violet: 1xCBS_Fw_Sox2P, green: 3xCBS_Fw_Sox2P, orange: 3xCBS_Rv_Sox2P, grey: wildetype (WT) non fluorescent control clone. Different linetypes indicate separately derived clones with the same reporter construct. **(D)** PCR genotyping of clones for successful RMCE. PCRs with genome and construct specific primers for the left and right side of the SB-genomic junction were performed and the resulting amplicons loaded on a 1% agarose gel. Clones with the same 3xCBS_Fw_Sox2P insert in LP-39 showed different reporter levels in Fig S3C, highlighted boxes I and II refer to one high and one low reporter expression clone. **(E)** Sanger sequencing of the highlighted bands in (D) confirmed the right insert in those highlighted clones. **(F)** Timecourse experiment to assess reporter expression of different clones with the same insert over 11 days. T1 reflects the first measurement day, T11 reflects measurement on day 11. Two clones containing the 3xCBS_Fw_Sox2P (clone 1 and 2, same as genotyped in S3D) and five clones containing 3xCBS_Rv_Sox2P construct were measured by FACS. On each day, a control cell line containing Sox2P in LP-39 kb was measured and normalized to. Reporter expression (mTurq) and *Sox2*::mCherry expression of all clones measured on the same day were normalized to the Sox2P reporter and *Sox2*::mCherry values in LP-39 kb. **(G)** Comparison of the reporter expression score derived from the hopping experiments to the clonal integrations at LP-116 kb and LP-39 kb per construct. Each red dot for the clones represents a measured clonal cell line. The vertical line represents the mean value. The single blue dot for the reporter expression score is the median bootstrapped value at the overlapping LP windows. Each experiment type was normalized to the values of Sox2P per launch pad.

**Fig S4.**
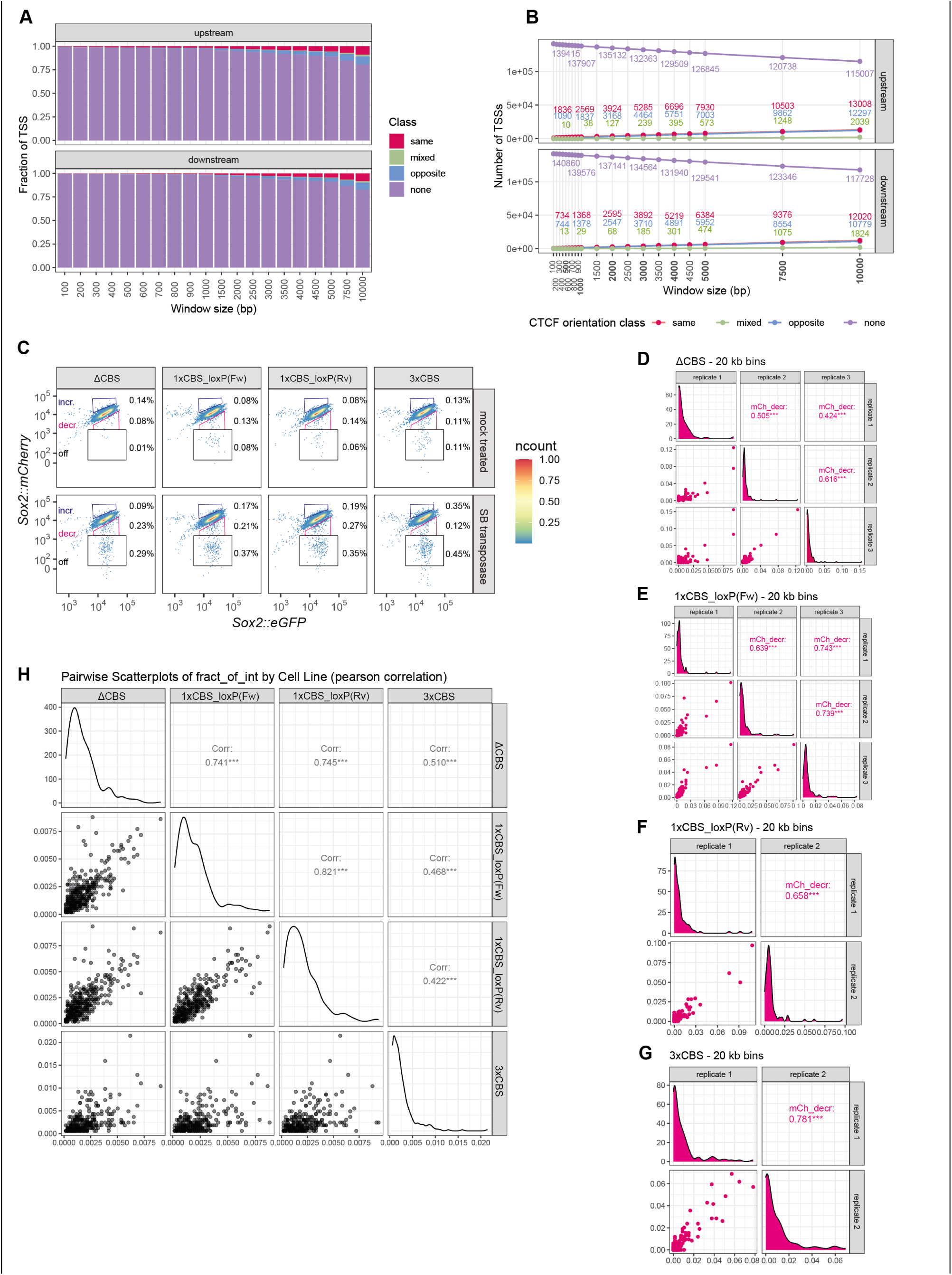
Genome wide CBS-TSS bias and reproducibility of CBS only hopping. **(A)** Fraction of TSSs in different window sizes (100–10000) that have no CBS, or are in the same, opposite or mixed orientation as the CBSs. Analysis was done for CBSs that are upstream or downstream of TSS. **(B)** Absolute number of differently classified TSSs for different window sizes containing the number of TSSs without any CBSs. **(C)** Expression levels of *Sox2*::mCherry (experimental) and *Sox2*::eGFP (control) in mock transfected and in cells transfected with SB transposase measured by FACS. The percentage of cells in each gate is indicated. Representative result of replicates. **(D-G)** Reproducibility of different SB construct integrations across the *Sox2* locus between biological replicates. Scatterplots show the fraction of population integration per 20kb bin (on a 2Mb interval as shown in Fig. S5A) colored by sorted population, across different replicates from LP-116 kb. Diagonal panels show distributions per replicate; upper panels show Spearman correlation coefficients. **(D)** shows correlation between three replicates of ΔCBS; **(E)** between three replicates of 1xCBS_loxP(Fw); **(F)** between two replicates of 1xCBS_loxP(Rv); **(G)** between two replicates of 3xCBS **(H)** Correlation of unsorted control integrations between the combined replicates of cell lines with different construct. Scatterplots show the fraction of unsorted integrations per 5kb bin (on a 2Mb interval, shown in Fig. S5A).

**Fig S5.**
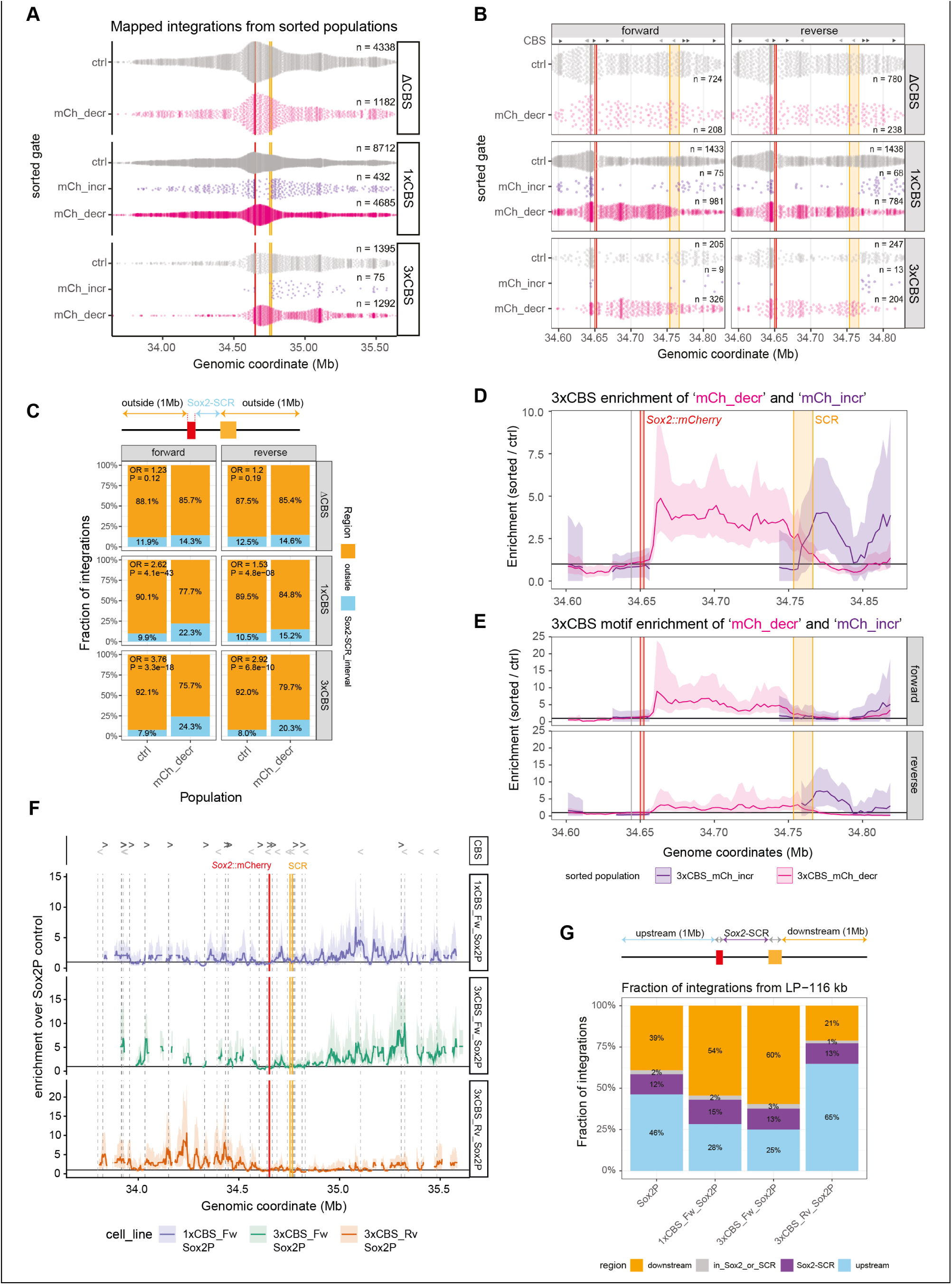
CBS integrations show motif specific effect on Sox2 expression. (**A**) Mapped integrations from unsorted (ctrl) and sorted cell populations (‘mCh_decr’ & ‘mCh_incr’) mobilized from LP-116 kb displayed per transposon insert (ΔCBS, 1xCBS, 3xCBS). Each dot indicates one integration. Number of integrations plotted in a 2Mb interval (chr3:33643960-35643960) is represented by n. **(B)** as in (A), except plotting range was changed to (chr3: 34590000-34830000). **(C)** Distribution of ΔCBS, 1xCBS and 3xCBS integrations hopped from LP-116 kb into two regions: (1) outside region, which are integrations mapping 1Mb upstream of the *Sox2*::mCherry gene and 1 Mb downstream of the start of the SCR. (2) Integrations spanning the *Sox2*-SCR interval. Mapped integrations were stratified by motif orientation and a Fisher exact test was performed for each motif orientation to examine significant differences between integrations outside vs inside in control vs ‘mCh_decr’ sorted population. P-value and odds ratio (OR) are reported. **(D)** Enrichment of 3xCBS integrations sorted for reduction (‘mCh_decr’) and increase in *Sox2*::mCherry (‘mCh_incr’) over the normal distribution of integrations (unsorted control). Enrichment was smoothened using a 25kb window, shifted by 5kb steps and only plotted if at least one integrations per window are present. Shaded region indicates 95% CI. **(E)** as in (D), except enrichment was calculated per motif orientation with a 25kb window, shifted by 5kb steps and only plotted if at least one integrations per window are present. **(F)** Enrichment of CBS containing reporter integrations (1xCBS_Fw_Sox2P, 3xCBS_Fw_Sox2P, 3xCBS_Rv_Sox2P) over the Sox2P control cell line was calculated using a smoothened 20kb window, shifted by 2kb steps and only plotted if a least three integrations per window were present. **(G)** Distribution of reporter integrations (Sox2P, 1xCBS_Fw_Sox2P, 3xCBS_Fw_Sox2P, 3xCBS_Rv_Sox2P) into four defined regions. (1) upstream: integrations that fall within 1Mb upstream LP-116kb; (2) downstream: integrations that fall within 1Mb downstream the end of the SCR; (3) *Sox2*-SCR: integrations between the end of *Sox2*::mCherry and the beginning of the SCR; in_Sox2_or_SCR: integrations directly in the *Sox2*::mCherry or SCR. The fractions of integrations into each of this four regions were calculated and displayed per reporter construct.

**Fig S6.**
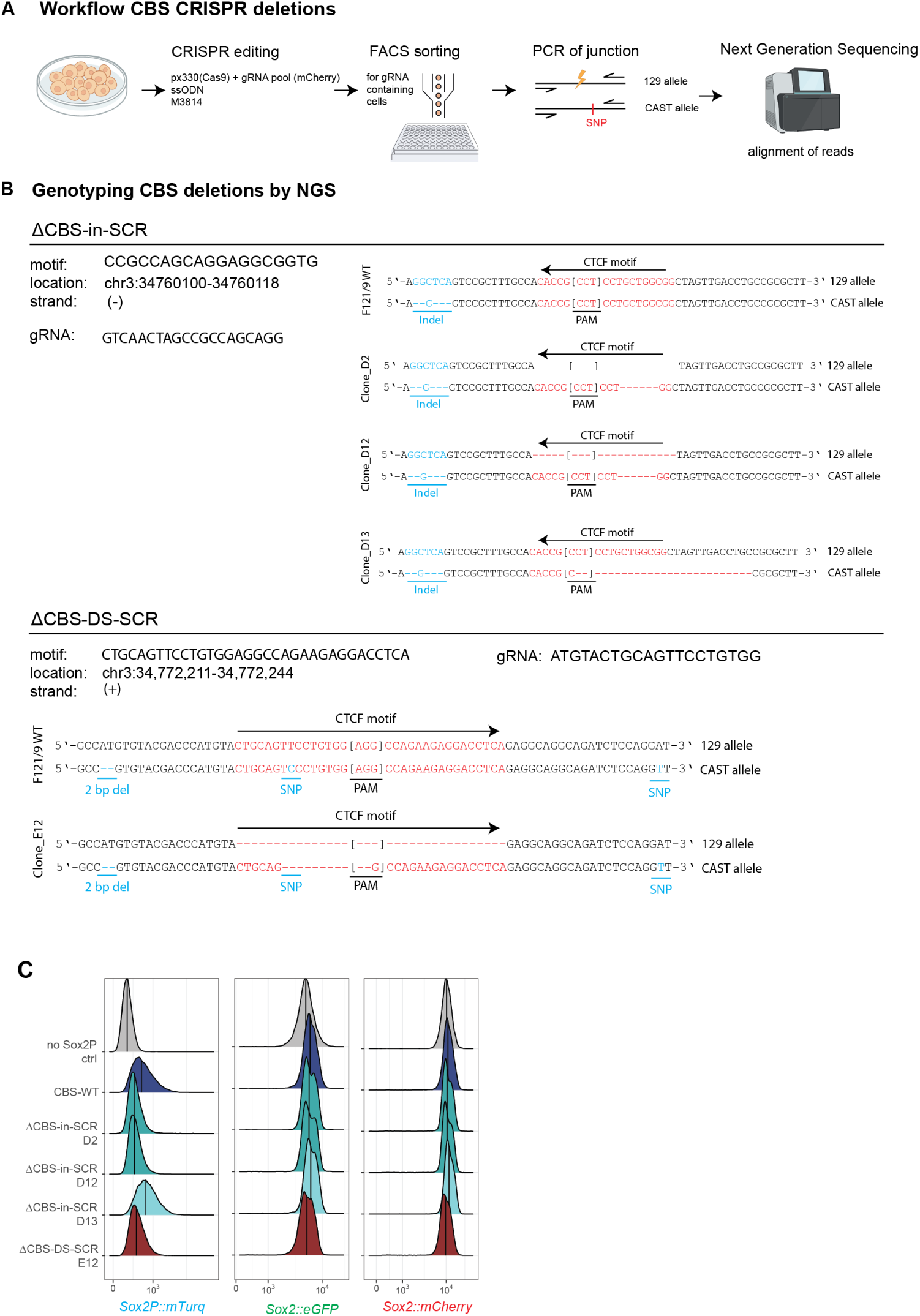
Generation and validation of endogenous CBS deletions in and downstream of the SCR. (**A**) Schematic workflow of using CRISPR/Cas9 do delete endogenous CBS. **(B)** Genotyping of selected CRISPR/Cas9 edited clonal cell lines. Validation of CBS deletions by next generation sequencing of ΔCBS-in-SCR and ΔCBS-DS-SCR clones. **(C)** Phenotyping endogenous *Sox2* (*Sox2*::mCherry, *Sox2*::eGFP) and reporter expression (Sox2P::mTurq) of control (no Sox2P WT ctrl & CBS-WT) and CBS deletions clones (ΔCBS-in-SCR clones D2, D12, D13 and ΔCBS-DS-SCR clone E12) by FACS.

**Fig S7.**
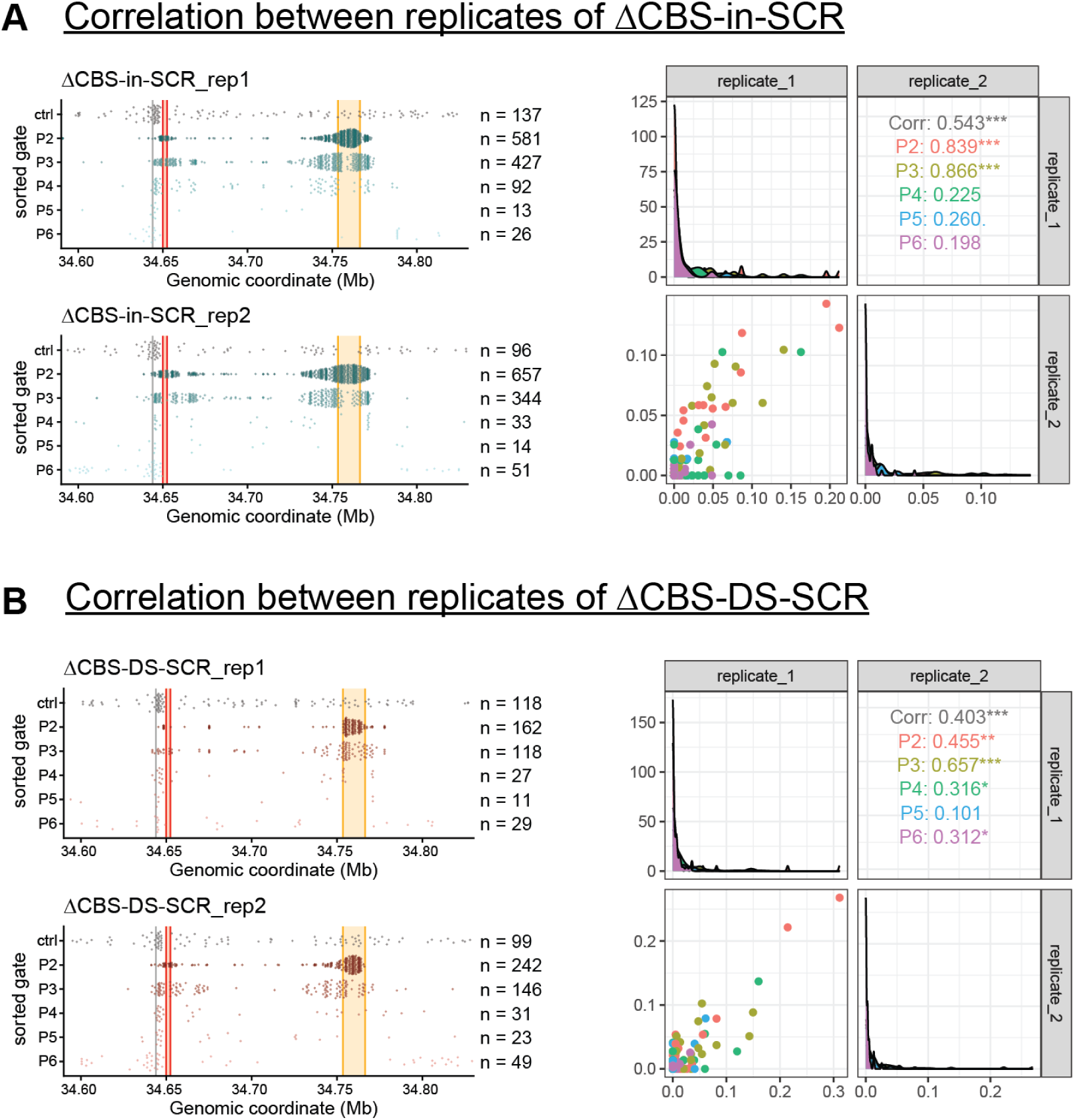
Reproducibility of Sox2P hopping in ΔCBS-in-SCR & ΔCBS-DS-SCR cell lines from LP-116 kb. (**A**) *Left:* Distribution of Sox2P integrations from LP-116 kb in ΔCBS-in-SCR deletion cell line of two biological replicates across the native *Sox2* locus. Each dot represents a mapped integration. Number of integrations plotted is represented by n. *Right:* Reproducibility of reporter integrations across the *Sox2* locus between biological replicates. Scatterplots show the fraction of population integration per 5kb bin (on the interval in Fig S7A, left) colored by sorted population, across two replicates from LP-116 kb. Diagonal panels show distributions per replicate; upper panels show Spearman correlation coefficients. **(B)** *Left*: same as (A)-Left, except for ΔCBS-DS-SCR deletion cell line. *Right*: same as (A)-Right, except for ΔCBS-DS-SCR deletion cell line.

## Acknowledgments

We thank Giorgetti lab for sharing the *Sox2* fosmid; the NKI Genomics, Flow Cytometry, Protein Production and Research High Performance Computing core facilities for excellent support and members from our lab and the divisions of Gene Regulation and Molecular Genetics for inspiring and helpful discussions.

## Author Contribution

Conceptualization: M.E., B.v.S.

Wet-lab experiments: M.E., C.J.I.M., M.K., L.B.

Data analysis and visualization: C.J.I.M., M.E.

Method development: M.E., C.J.I.M.

Code development: C.J.I.M., M.E.

Writing: M.E., C.J.I.M., B.v.S.

Funding Acquisition: B.v.S., C.J.I.M.

Supervision: B.v.S., E.d.W.

## Data and materials availability

Laboratory notebooks and supplementary data (including plasmid sequences, FACS data, etc.) are available on Zenodo (10.5281/zenodo.18955913). Raw sequencing data is available on GEO under accession numbers GSE324456.

Code and supplementary files used as input for the scripts are available on Zenodo (10.5281/zenodo.18955085) and GitHub. All plasmids used will be made available upon request.

## Material and Methods

### Cell culture

F121/9 (CAST/EiJ × S129/Sv) female mouse embryonic stem cells (mESCs; RRID:CVCL_VC42) (*40*) and all derived clones were maintained in Serum+LIF+2i medium, as previously described (*23*). Cells were cultured on 0.1% gelatin-coated tissue culture plates in Glasgow minimum essential medium (GMEM; Sigma-Aldrich, G5154) supplemented with 15% fetal bovine serum (Thermo Fisher Scientific, 10270-106), 1% L-glutamine (Thermo Fisher Scientific, 25030024), 1% sodium pyruvate (Thermo Fisher Scientific, 11360-70), 1% MEM non-essential amino acids (Thermo Fisher Scientific, 11140-50), 1% penicillin–streptomycin (Thermo Fisher Scientific, 15070063), 100 µM β-mercaptoethanol, 10 x 4 U leukemia inhibitory factor (LIF; esg1107, Millipore), 1 µM MEK inhibitor PD0325901 (Mirdametinib, MedChemExpress), and 3 µM GSK-3β inhibitor CHIR99021 (Laduviglusib, MedChemExpress). Cells were grown at 37 °C in a humidified incubator with 5% CO₂ and passaged every 2 days. Mycoplasma contamination was monitored and excluded by regular testing (Lonza, LT07-318).

### eGFP/mCherry-tagged *Sox2* mESC line

The allele-specific *Sox2* reporter mESC line used in this study was generated previously from the F121/9 (CAST/EiJ × 129S1/Sv) background by CRISPR/Cas9-mediated homology-directed repair, as described before (*23*).

### Genetic engineering of the *Sox2* locus: Establishing launch pads LP-116 kb & LP-39 kb

The launch pad cell lines LP-116 kb and LP-39 kb generated previously (*23*) were used as founder cell lines. Landing pad cassettes were introduced into the *Sox2* locus of the *Sox2*-eGFP/mCherry F121/9 mESC line as described previously (*23*).

### RCMC donor plasmids

#### Sox2 reporter construct (Sox2P)

The *Sox2* promoter reporter cassette used for RMCE has been described previously (*23*). Briefly, a 2,346 bp fragment of the mouse *Sox2* promoter including the 5′ UTR (fosmid WL1-1017O07; mm9 chr3:34535173–34570689, kindly provided by Luca Giorgetti) was PCR-amplified with Phusion polymerase using GC-rich buffer and DMSO, and subcloned into the StuI-digested pCR™-Zero backbone (Addgene #120275). The *Sox2* promoter fragment was then excised with XbaI and SalI and cloned upstream of mTurquoise2 (Addgene #118617) to generate a Sox2P–mTurquoise2 reporter, followed by insertion of an SV40 polyadenylation signal 3′ of the fluorophore. Finally, heterotypic F3 and FRT sites were added flanking the cassette by PCR to generate the RMCE donor plasmid pME034 (sequence in **Table S5**). All intermediate and final constructs were validated by Sanger sequencing; primer sequences are listed in **Table S1.**

#### 1xCBS-Fw-Sox2P construct

To generate the 1xCBS-Fw-Sox2P construct, the Sox2P reporter plasmid pME034 was digested with BamHI and AcII, and the linearized backbone was gel-purified. As BamHI cuts upstream of the FRT sequence in pME034, we designed a long forward primer (MEP219) containing a BamHI site, an FRT site, and a single CTCF binding site (1×CBS), and used it together with a reverse primer annealing at the AcII site in pME034 (MEP219) to amplify the corresponding fragment. The PCR product was subcloned into StuI-digested pCR™-Zero (Addgene #120275), and colonies were screened by PCR to identify clones with the correct insert. Plasmids confirmed by colony PCR were digested with BamHI and AcII, and the resulting 575 bp FRT–1×CBS-Fw fragment was gel-purified. The BamHI/AcII-digested pME034 backbone and the FRT–1×CBS-Fw fragment were ligated using the Quick Ligation kit (NEB, M2200S) at a 1:4 vector:insert ratio to generate plasmid pMK03 (FRT–1×CBS-Fw–Sox2P–mTurquoise2–SV40 polyA–F3). The final construct was validated by Sanger sequencing. Primer sequences are provided in **Table S1** and RMCE sequence is provided in **Table S5.**

#### 3xCBS-Fw & 3xCBS-Rv construct

The 3xCBS construct was based on previously published sequences (*16, 18*). The 3xCBS-Fw and 3xCBS-Rv sequences were flanked by FRT and F3 sites and ordered as clonal genes (plasmids) via Twist Bioscience. We re-transformed and verified plasmids (pME038 & pME039) by Sanger sequencing. Sequences of 3xCBS-Fw and 3xCBS-Rv can be found in **Table S5.**

#### 3xCBS-Fw-Sox2P & 3xCBS-Rv-Sox2P construct

To generate the 3xCBS-Fw-Sox2P & 3xCBS-Rv-Sox2P constructs, we PCR amplified the Sox2P-mTurq-polyA region from pME025 with primers containing compatible Gibson assembly overhangs (MEP215 + MEP216). Next, we digested the plasmid containing the FRT-3xCBS-Fw-F3 (pME038) or FRT-3xCBS-Rv-F3 (pME039) array with SpeI-HF (NEB, R3133S) and purified the linearized plasmid via gel extraction. We assembled the Sox2P-mTurq-polyA fragments together with the linearized plasmid with NEBuilder® HiFi DNA Assembly Master Mix (NEB, E2621S) to create the final plasmids containing FRT-3xCBS(Fw)-Sox2P-mTurq-polyA-F3 called pMK01 and FRT-3xCBS(Rv)-Sox2P-mTurq-polyA-F3 called pMK02. The final constructs were validated by Sanger sequencing. Primer sequences are provided in **Table S1**. RMCE insert sequences are provided in **Table S5.**

#### 1xCBS-Rv-loxP(Fw) & 1xCBS-Rv-loxP(Rv)

The plasmids (lb002-fw & lb002-rv) containing the 1xCBS reverse motif (chr2:48886253-48886271; ATACTGCCCTCTGCTGTTCA) flanked by forward (lb002-fw) or reverse (lb002-rv) oriented loxP sites were a gift from the de Wit lab. The plasmids were digested with SpeI-HF (NEB, R3133S) and NheI-HF (NEB, R3131S) and cloned in between Sleeping Beauty arms of a target vector. The inserts were PCR amplified with primers containing FRT (MEP74) and F3 sites (MEP131) using NEBNext High-Fidelity 2x PCR Master Mix (NEB, M0544) and cloned into StuI digested pCR™-Zero backbone (Addgene 120275) using the Takara ligation mix according to the manufacturers protocol. Five μL of ligation mix were transformed into 100 uL of DH5α (NEB, C2987H) chemically competent cells. Colonies were screened by colony PCR and validated by Sanger sequencing. The final plasmids were named pME013 (FRT-SB-lb02_loxP_Fw-SB-F3) and pME016 (FRT-SB-lb02_loxP_Rv-SB-F3). Insert sequences are provided in **Table S5.**

### Recombination mediated cassette exchange (RMCE)

#### Integration into launch pads (LP’s –116 kb and –39 kb)

Reporter and control constructs were loaded into the Sleeping Beauty (SB) launch pad cassette by recombination-mediated cassette exchange (RMCE), as described previously (*23*). Briefly, 3 × 10^5^ cells were transfected with 0.5 µg of a Flp recombinase–encoding plasmid (Addgene #13787) and 2 µg of donor plasmid using 7.5 µL Lipofectamine 2000 (Invitrogen, 11668019). Two days after transfection, cells were seeded sparsely and ganciclovir (2.5 µg/mL) was added to the medium to selectively eliminate HyTK-expressing cells. Ganciclovir-resistant colonies were picked, expanded, and screened by PCR to identify correctly recombined clones.

### Establishing ΔCBS cell line

The 1xCBS-Rv-loxP(Fw) construct was integrated in LP-116 kb of *Sox2* double tagged mESCs via RMCE. After selection with ganciclovir, clones were grown and picked for genotyping. Three clones were validated by PCR amplification and subsequent Sanger sequencing (subclone 1, 12, 16). Subclone 16, containing the 1xCBS-Rv-loxP(Fw) construct in LP-116 kb, was used to generate the ΔCBS cell line. Cells were seeded and transfected with a plasmid expressing CRE-Puro (pJK14_946_CRE_Puro) with subsequent selection at 1 ug/ml Puromycin. Surviving cells were seeded as single cells in a limited dilution and grown out. Six clones were picked and successfully genotyped by PCR for the deletion of the CBS. Furthermore cells were checked for SOX2 expression by FACS. The resulting cell line is referred to as ΔCBS.

### Establishing ΔCBS-in-SCR and ΔCBS-DS-SCR cell lines

To delete the ‘ΔCBS-in-SCR’, we ordered a gRNA previously published by another group (*11*) together with a ssODN repair oligo. For the ‘ΔCBS-DS-SCR’, we designed five gRNAs targeting the CBS region together with a ssODN repair template. gRNAs were cloned into an expression plasmid containing mCherry. One guide per region was selected (pCM036 for ‘ΔCBS-in-SCR’ and pCM032 for ‘ΔCBS-DS-SCR) and 0.5 million cells were transfected with 1 ug Cas9 plasmid (pX458, Addgene 48138) and 1.5 ug gRNA plasmid using 7.5 ul lipofectamine 2000. Additionally, for ‘ΔCBS-in-SCR’, we used 1.5 ul of a 10 uM working stock of ssODN008, for ‘ΔCBS-DS-SCR’ we used 1.5 ul of a 10 uM working stock of ssODN007. Both transfections were carried out in the presence of an DNAPKcs inhibitor, namely M3814 (MCE, HY-101570). The cells were cultured in the presence of the inhibitor for two days and subsequently sorted for eGFP/mCherry expression. We verified the allele specific CBS deletions by high-throughput sequencing **(Sup Fig. S6A & S6B)** amplifying the ‘ΔCBS-in-SCR’ region (CMP095+CMP096) and the ‘ΔCBS-DS-SCR’ region (CMP097+CMP098) with locus specific primers for 15 cycles. PCR2 was performed for another 15 cycles with TAC0159.x and TAC0009, allowing for different index use and sample multiplexing. Sequences of gRNAs and ssODN’s used in this study can be found in **Table S2** and **Table S3**.

### SB hopping

For each cell line and biological replicate, Sleeping Beauty (SB) transposition was induced by transient expression of SB100X (pME07). Specifically, 2.5–5 × 10^6^ cells were transfected with the bicistronic plasmid pME07, which encodes the SB100X transposase and the human low-affinity nerve growth factor receptor (LNGFR), using 5 µg plasmid per 10^6^ cells and 15 µL Lipofectamine 2000 (Invitrogen, 11668019). As a negative control for SB-mediated hopping, 5 × 10^5^ cells were transfected with a similar bicistronic plasmid encoding PiggyBac (PB) transposase and LNGFR (pLD042). Thirty hours after transfection, LNGFR-positive cells were enriched by magnetic cell sorting (MACS) using MS columns (130-042-201, Miltenyi Biotec) and LNGFR MicroBeads (130-091-330, Miltenyi Biotec), according to the manufacturer’s instructions.

#### Hopped cell pools

One week after transfection, 2 × 10^5^ SB-transposase-transfected cells were set apart as an unsorted pool and the remaining cells were sorted by FACS (BD FACSAriaTM Fusion Flow Cytometer) into pools for populations based on mTurq expression level, using the following lasers and filters: mTurq: 442nm V[F] 470/20, eGFP: 488nm BL[B] 530/30, mCherry: 561nm YG[D] 610/20 (see **Fig. 2B** for the exact gates). PiggyBac (PB) transposase transfected cells served as mock-treated FACS control. In the first biological replicate of the –116 kb Sox2P-reporter (**Fig. S2B, rep1**), cells were only selected based on mTurq level, in all later replicates we selected only *Sox2::mCherry*-positive cells for all *Sox2::mCherry* containing cell lines. To increase the number of integrations, we also sorted one pool of 50,000 cells for one replicate of – 116kb_Sox2P population P3-P5. For CBS only constructs, *Sox2*::mCherry expression was gated into three gates, whereas the lower gate indicates deletions (**Fig. 5C**). Sorted pools were expanded and crude lysates were obtained from a full 96 well plate by lysing with 50 µL DirectPCR Lysis Reagent (102-T, Viagen Biotec) supplemented with 100 µg/mL proteinase K, and incubating 2.5 h at 55 °C and 45 min at 85 °C. For the 50,000 cell pools and the unsorted cell pools, gDNA was extracted after expansion using the ISOLATE II Genomic DNA Kit (BIO-52067, Bioline).

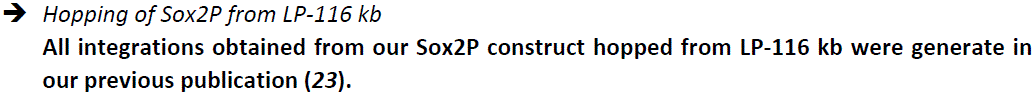

#### Establishing clones

To generate clonal cell lines from 1xCBS-Fw-Sox2P constructs, we FACS sorted single cells with P1 reporter expression and grew them out. After expansion, crude lysates were obtained from full 96-well plates as discussed above. Mapping was done via Tn5-tagmentation of the crude lysates.

### Tagmentation-based mapping of SB integrations

Sleeping Beauty integration sites were mapped by tagmentation-based amplification and sequencing of ITR–genome junctions, as described previously (*23*), corresponding to (*41, 42*). In brief, tagmentation was performed on either 100 ng genomic DNA or 3 µL crude lysate, followed by linear enrichment of ITR–genome junctions using SB-ITR–specific primers (MEP009 for the 5′ ITR, LD027 for the 3′ ITR) and a first PCR (PCR1) with primers MEP011 (5′ ITR) or MEP034 (3′ ITR). For clonal samples and sorted pools, multiple parallel tagmentation reactions were performed to increase integration-mapping depth. Libraries were sequenced with 150 bp paired-end reads on Illumina MiSeq or NextSeq 550 instruments with a 10% PhiX spike-in.

### Flow Cytometry

Allele-specific SOX2 expression and reporter fluorescence of clones and pools were measured on a BD LSRFortessa™ equipped with a High Throughput Sampler (HTS) for 96-well plate acquisition. Fluorescence was detected using the following laser/filter settings: eGFP (488 nm, 530/30), mCherry (561 nm, 610/20), and mTurquoise2 (405 nm, 470/20). Single-color controls confirmed no spillover of mTurquoise2 into the GFP channel. Single cells were gated in FlowJo, and fluorescence data were exported as.fcs files for analysis in R using the flowCore package (*43*).

Cell sorting was performed on a BD FACSAria™ Fusion using the same filter configuration, but mTurquoise2 was excited with a 445 nm laser for improved performance. To correct for spectral overlap, 1.6% of the 470/20 signal (mTurquoise2) was subtracted from the 530/30 GFP channel. Automatic background signal subtraction resulting from the fluid between cells is applied by the FACS machine.

## Computational analysis

### Tagmentation mapping

We used the same pipeline as published earlier (*23*)to map SB integrations. Briefly, reads were mapped to the mouse reference genome (Release M23 – mm10) and filtered for a minimal mapping quality of 10. For the sorted pools, integrations with at least two unique mapping reads were retained. To ensure that high reporter expression is due to location and not due to hopping-induced *Sox2*::mCherry deletions, the integrations in the P1 pools were further filtered for having at least one read mapping each from the 5′ ITR and 3’ ITR (reverse and forward read). For the large unsorted control pools we filtered integrations less stringently, including any integration with at least one mapping read. Note that the integrations in these large unsorted pools are not mapped exhaustively. In a few experiments, integrations exactly matching the –39 kb launch site (chr3:34721183-34721192) were found across many pools. This indicated a PCR contamination from an earlier experiment. Therefore in each experiment we filtered out all integrations exactly matching those coordinates with the exception of the launch pad that the reporter was mobilized from. Furthermore, we filtered out 18 overrepresented integrations (most likely bleedtrough sequencing contamination from other clonal cell lines) present in all three cell lines of 1xCBS_CBS_Fw_Sox2P, 3xCBS_Fw_Sox2P & 3xCBS_Rv_Sox2P of replicate 2.

#### Insertion clones

To identify the true integrations in clonal cell lines, we filtered more stringently, requiring at least ten unique mapping reads. Next, we discarded any locations that were supported by <20% of the read count of the top location, since these are likely to be either index-swapped reads from other clones or off-target reads. To call an insertion with confidence, we required to find exactly one integration supported by 5’ ITR and 3’ ITR mapping. Any clones with more mapped locations were manually reviewed: any clones for which the extra reads could be explained by index-swapping were included, while clones with evidence of a secondary integration or an unexplained high number of mapping reads from a secondary location were discarded.

### Genotyping of CBS-deletion clones by HTS

To verify the allele-specific CBS deletions in the high-throughput sequencing data, the reads were first pre-processed by trimming the amplification primers and Illumina adaptors with a modified version of cutadapt {Martin, 2011, https://doi.org/10.14806/ej.17.1.200}. The first 128 (ΔCBS-in-SCR) or 186 (ΔCBS-DS-SCR) bp of each read were extracted and clustered using starcode with Levenshtein distance 0, to count identical reads (*44*). For each clone, the three most abundant reads were written into a fasta file, aligned to the amplicon sequence from mm10 in SnapGene and manually compared to the annotated indels and single nucleotide polymorphisms from the 129S1 and CAST allele from the Mouse Genomes Project (https://www.mousegenomes.org/publications/, (*45*) (*46*) (**Fig. S6B**). Clones were only one allele was detected were excluded. Then clones were selected that have a deletion of the CTCF motif on the 129S1 allele, regardless of the state of the CAST allele. A clone with a deletion on the CAST allele and an unedited sequence on the 129S1 allele was used as control (ΔCBS-in-SCR D13, **Fig. S6C**)

### Expression score calculation or reporter integrations

The expression score was calculated the same way as previously described (*23*) “To calculate expression scores from sorted pools, we first combined the integrations from each experiment per sorted population. Identical integrations found in multiple sorted pools from one experiment most likely originated from a single hopping event. Still, we treated them as separate data points because they were independently sorted into an expression gate. Next, we estimated the fluorescence intensity of the hopped reporters in each expression gate (*Fluo_p_*) from the median fluorescence intensity as measured during the FACS sorting process.

Between different experiments and populations, a different number of cells was sorted. These sorted cells (and thus the mapped integrations) also represent a different fraction of the unsorted cell population (see supplementary note 1). To account for this, we determined for each population *P* the correction factor *Fsort_p_* (using a pseudo-count of 1 to account for sorted populations that were too rare to observe in the FACS recording):

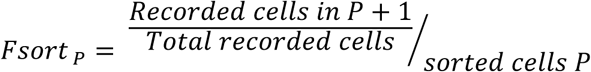

*Fsort* can be interpreted as the fraction of the original unsorted cell population that is represented by each sorted cell or mapped integration (under the assumption that each sorted integration is equally likely to be mapped). When for one cell line the number of replicate experiments was unequal between different populations, this was also accounted for in *Fsort_p_* by multiplying by

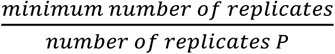

Next, for any given genomic window (W), the expression score was defined as the average of the population fluorescence (Fluo_P_), weighted by the number of integrations of each population in that window times the correction factor *Fsort_p_*.

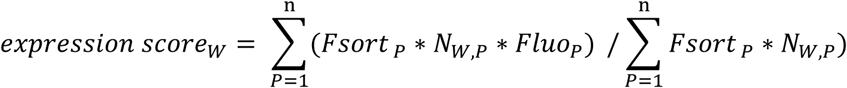

When pooling replicate data, *Fsort_p_* and *Fluo_p_* could vary between replicates. Therefore, the integrations were counted separately per replicate. When computing the Sox2P reporter expression score based on the three launch pads combined, their replicate experiments were combined in a similar way (6 replicates total: 2 replicates from –161kb, 3 replicates from –116kb [see supplementary note], 1 replicate from +715kb):

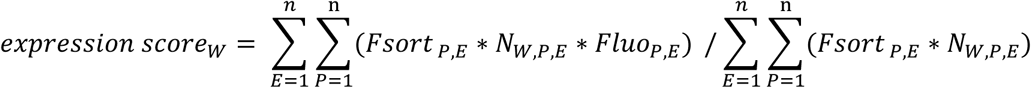

To obtain a running window expression score (e.g. 10 kb window), integrations were counted in smaller bins (e.g. 1 kb, this is the step size) and the expression score was calculated from a group of bins (e.g. 10 bins). Window and step size were selected based on the density of integrations in each experiment and region of interest. To account for the deletion in the ΔCBS_*Sox2::mCherry* cell line, we shifted all integrations after the deletion by the width of the deletion, calculated the running mean expression score and then shifted the bins after the deletion back to actual their genomic position. To estimate a 95% confidence interval, the genome-wide list of integrations for each cell line was bootstrapped 1000 times and the expression score was calculated from each bootstrap. Reported are the median of these bootstrapped expression scores as well as the 2.5^th^ to 97.5^th^ percentile (95% confidence interval), for each window with at least 3 sorted integrations. To obtain the strand-specific expression score, only integrations on a specific strand were included in integration count of a window (*N_W,P,E_*).”

Based on the calculation of the expression score, the hopping induced bias of different inserts **(Fig. 6B & 6C)** does not skew the calculation of the levels of the expression score.

### Calculation of Enrichment (Fig. 5D-H, S5D, S5E)

To quantify how frequently integrations sorted for a phenotype *P* (e.g. “mCh_decr”) are found at a genomic position relative to the non-sorted control population (“ctrl”) of the same cell line, we computed the enrichment in a sliding window.

For any given genomic window (W), we first computed the fraction of sorted integrations in that window over the total mapped sorted integrations genome-wide (*fract_W,P_*).

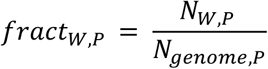

Then we calculate the same fraction for the control integrations (*fract_w,ctrl_*). The enrichment of sorted integrations over control integrations in the window W is then defined as:

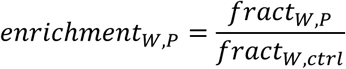

To obtain the enrichment in a sliding window (e.g. 10 kb window), integrations were counted in smaller bins (e.g. 1 kb, the step size) and then the enrichment was calculated from the summed integrations from a group of bins (e.g. 10 bins).

For motif orientation-specific analysis, integrations in a window was calculated for the forward and reverse motif integrations separately, as for instance:

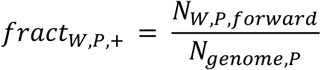

To estimate a 95% confidence interval, integrations from all populations (control and sorted) of the same cell line were resampled together with replacement (1000 times), and enrichment was recalculated from each bootstrap. Reported are the median of these boostrapped enrichment values as well as the 2.5^th^ to 97.5^th^ percentile (95% confidence interval), for each window with at least 3 integrations in the control for non-stranded analysis and 2 integrations for stranded analysis.

### Computational analysis Flowcytometry

To visualise the exact FACS gating used to sort SB-transposase–transfected cells **(Fig. 2B, 5C, 7C, S4C)**, we imported the FACSDiva experiment file (.xml) together with the raw FCS files into R using diva_to_gatingset from the CytoML package (*47*). Colour density plots were generated with ggcyto in combination with geom_hex.

To compare the *Sox2* and reporter expression levels between clones, the fluorescent values were corrected and normalized. First, the raw FCS files were gated for single cells in FlowJo and loaded into R with the flowCore package (*43*). Next, the median fluorescence intensity (MFI) was calculated for each fluorophore and each clone. The autofluorescence obtained from cells without fluorophore was subtracted from the respective MFIs of each sample. Subsequently, the corrected mCherry and mTurquoise2 values were normalized by the corrected eGFP value of the same sample, to adjust for any global effects on *Sox2* expression. Normalized mTurquoise2 and mCherry values of clones with the reporter integration at the same genomic position were averaged.

Because there was variation in the absolute MFIs between measurement days, we divided the normalized fluorescence values by the respective value of the original LP-116kb_Sox2P cell line, measured on the same day. In this way, we obtained relative *Sox2::mCherry* and reporter (mTurquoise2) expression levels that could be compared between measurement days and experiments (**Fig. 3B, 3C**). For the Timecourse measurements, each obtained clonal measurement was normalized to the respective measurement of the Sox2P insert in LP-39kb of the same timepoint, therefore correcting for day specific changes in FACS acquisition **(Fig. S3F)**.

When measuring the 1xCBS_Fw_Sox2P clones **(Fig. 3D)**, we didn’t measure a non-fluorescent wildtype control to estimate the autofluorescence. Therefore we took the average of two WT measurements from another experiment, measured on another day but on the same machine with the same laser settings and experimental layout.

### Comparison between expression score and clonal measurements

The median bootstrapped expression score (stranded) of the different inserts; Sox2P, 1xCBS_Fw_Sox2P, 3xCBS_Fw_Sox2P, 3xCBS_Rv_Sox2P; were compared to the relative reporter expression of the measured clones derived for LP-116 kb and –39 kb. Reporter expression and expression scores from different inserts were normalized to the Sox2P insert for their respective launch pad position (either LP-116 kb or LP-39 kb) and result in the relative reporter expression **(Fig. S3G)**.

### Genome wide CBS orientation analysis

To assess potential CTCF binding site biases (CBSs) genome wide, we analysed orientation biases of CBSs in regards to the orientation of TSSs. Positions of TSSs were obtained from GENCODE and used as strand-aware reference points. We used the AH104755 dataset from AnnotationHub containing genomic coordinates of FIMO-predicted CTCF binding sites for the mouse mm10 genome assembly. We filtered the putative CBSs for a p-value < 1e-6 which should result in a true positive rate of approximately 80% (*48*) {Hahne, 2009 #90}. For each TSS, upstream and downstream windows of increasing size (100 bp to 10 kb) were defined relative to transcriptional direction. Within each window, CTCF sites were classified into four mutually exclusive categories: same (all motifs oriented toward the TSS), opposite (all motifs oriented away from the TSS), mixed (both orientation present) and none (no CBS present in window). This classification was performed separately for upstream and downstream regions. For each window size, the fraction and absolute number of TSSs associated with each class were calculated genome-wide. In addition, the density of CTCF sites around TSSs (±10 kb) was computed separately for motifs oriented in the same or opposite direction from the TSS. Density profiles were plotted with a bandwidth of 250, strand-aligned and smoothed for visualization.

### Statistical analysis of integration biases

To statistically quantify differences in genomic integration patterns between cells sorted for a phenotype and their respective control, we assigned two genomic regions and calculated the fraction of integrations in each. *Region outside*: 1Mb upstream the *Sox2* gene (chr3: 33649995-34649995) & 1Mb downstream of the start of the enhancer (chr3:34753416-35766401); *Region Sox2-SCR*: chr3: 34652461-34753415. We used a Fisher Exact test to calculate the p-value and odds ratio (OR). We performed this analysis regardless of motif orientation **(Fig. 5E)** and with motif orientation **(Fig. S5C)**.

To assess if there is a significant hopping bias of integrations based on the transposon insert, we used a Fisher Exact test to compare integrations that are 1Mb upstream (chr3:33643960-34643960) or downstream (chr3:34643960-35643960) of LP-116 kb between Sox2P (ctrl) and different Sox2P containing inserts (1xCBS_Fw_Sox2P; 3xCBS_Fw_Sox2P; 3xCBS_Rv_Sox2P) as well as ΔCBS (ctrl) with 1xCBS_Rv and 3xCBS_Fw **(Fig. 6D)**.

### Public ChIP-seq datatracks

We searched the Cistrome Data Browser (*49*) for ChIP-seq datasets of P300, H3K27ac, SOX2, POU5F1 (OCT4) and NANOG obtained in mouse embryonic stem cells (filtering for the cell type “Embryonic Stem Cell’). Only datasets from non-perturbed or control conditions were selected. Next, the reprocessed bigwig tracks were downloaded from cistrome (http://dc2.cistrome.org//genome_browser/bw/X, where X is the cistrome ID) and visualized using the BigwigTrack function from Signac (*50*), smoothing over 100bp. References to the external datasets used can be found in **Table S6.**

**Table S1.**
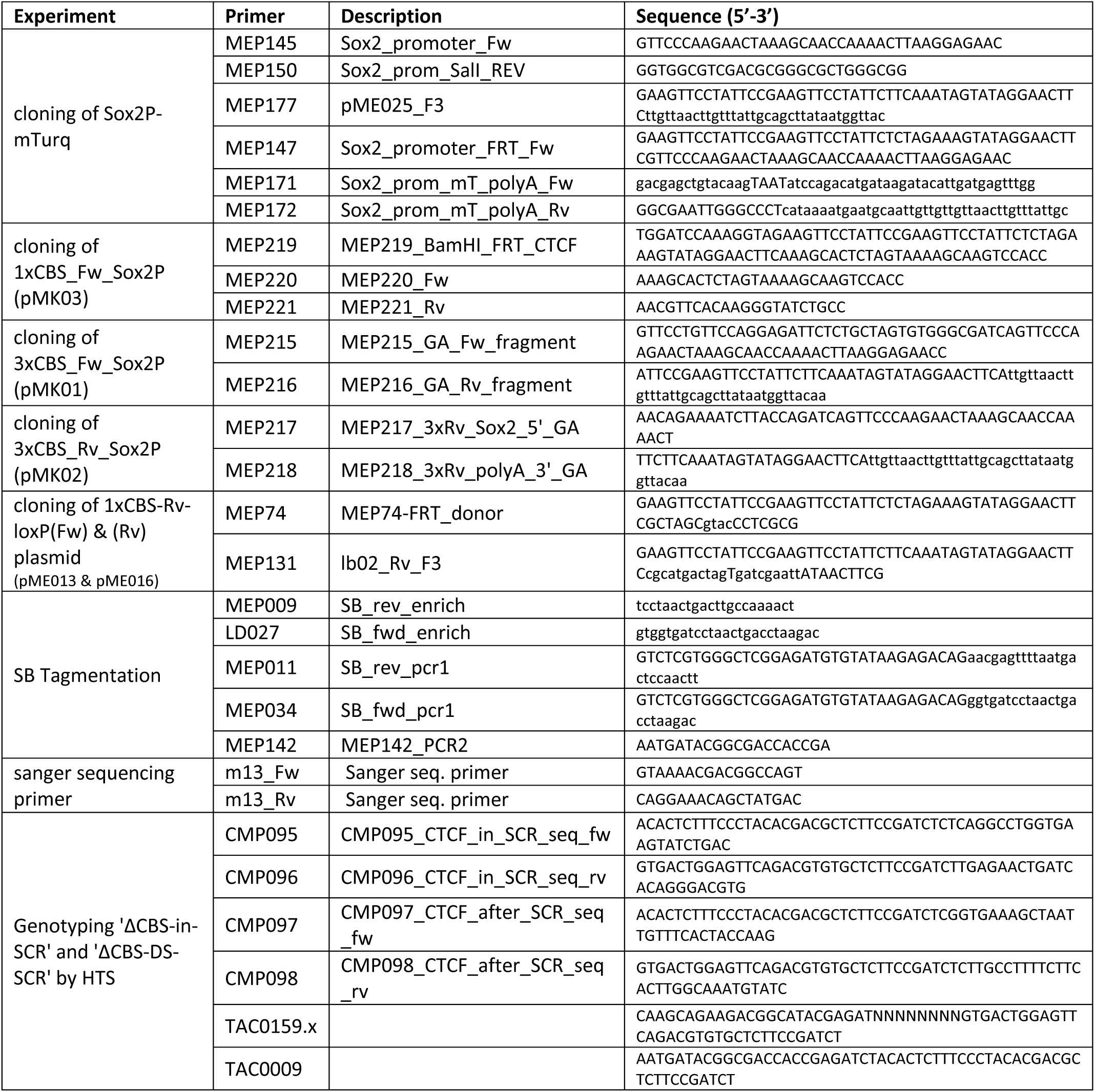
List of primers used in this study.

**Table S2.**
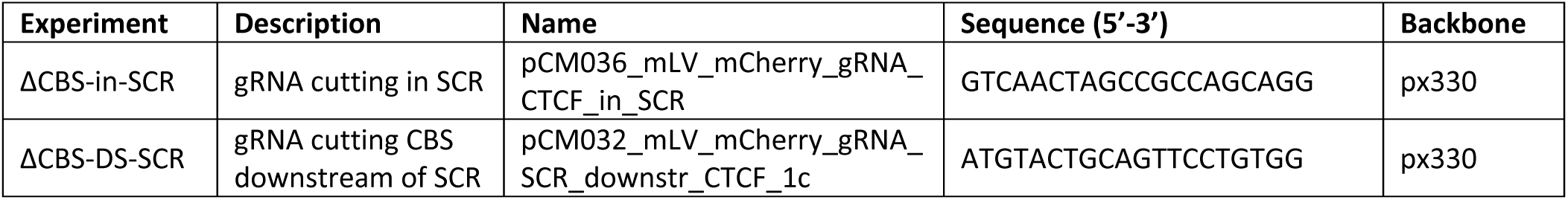
List of gRNAs used.

**Table S3.**
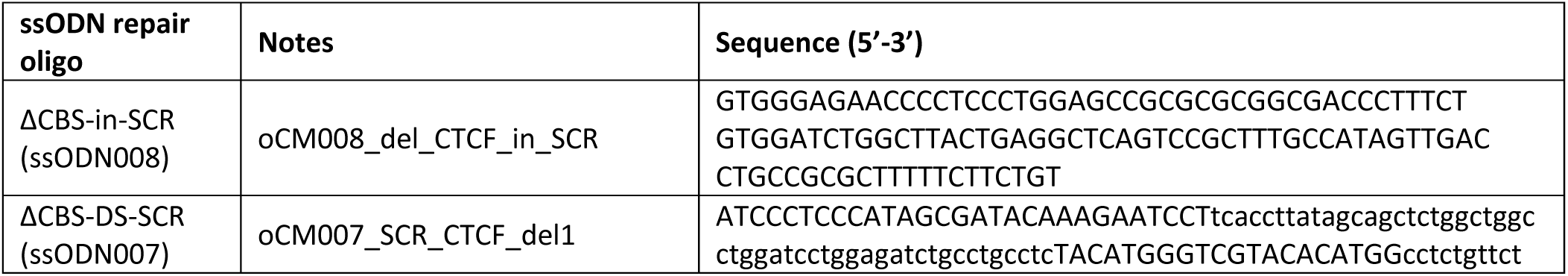
List of ssODN repair oligos used.

**Table S4.** List of plasmids used in this study.

| Plasmid name | Description | Addgene number (if available) |
| --- | --- | --- |
| pJK14 | pJK14 946 cre puro | X |
| pLD042 | pLD042_hPGK-PB-IRES-NGFR-polyA | X |
| pME07 | pME07_hPGK-SB100-IRES-NGFR-polyA | X |
| pME013 | pME013_lb02_Fw_pZero | X |
| pME016 | pME016_lb02_Rv_pZero | X |
| pME015 | pME015_SB-FRT-mPGK-HyTK-F3-SB | X |
| pME024 | pME024_HA-SB-HyTK-HA_Sox2 gRNA#8_KnockIN construct | X |
| pME034 | pME034_FRT_Sox2Prom_mTurq2_polyA_F3_pZero | X |
| pME038 | pME038_3xCTCF_Fw_motif_FRT_F3 | X |
| pMK01 | pMK01_3xCTCF_Fw_Sox2P_mTurq | X |
| pMK02 | pMK02_3xCTCF_Rv_Sox2P_mTurq | X |
| pMK03 | pMK03_Endogenous_CTCF_Fw_Sox2P_mTurq | X |
| pCM032 | pCM032_mLV_mCherry_gRNA_SCR_downstr_CTCF_1c | X |
| pCM036 | pCM036_mLV_mCherry_gRNA_CTCF_in_SCR | X |
| pX330-addgene | pX330-gRNA | 158973 |
| pCR-Zero | pCR Zero | 120275 |
| pCAG-Flpe-addgene | pCAG-Flpe_addgene-plasmid-13787-sequence-7769 | 13787 |

**Table S5.**
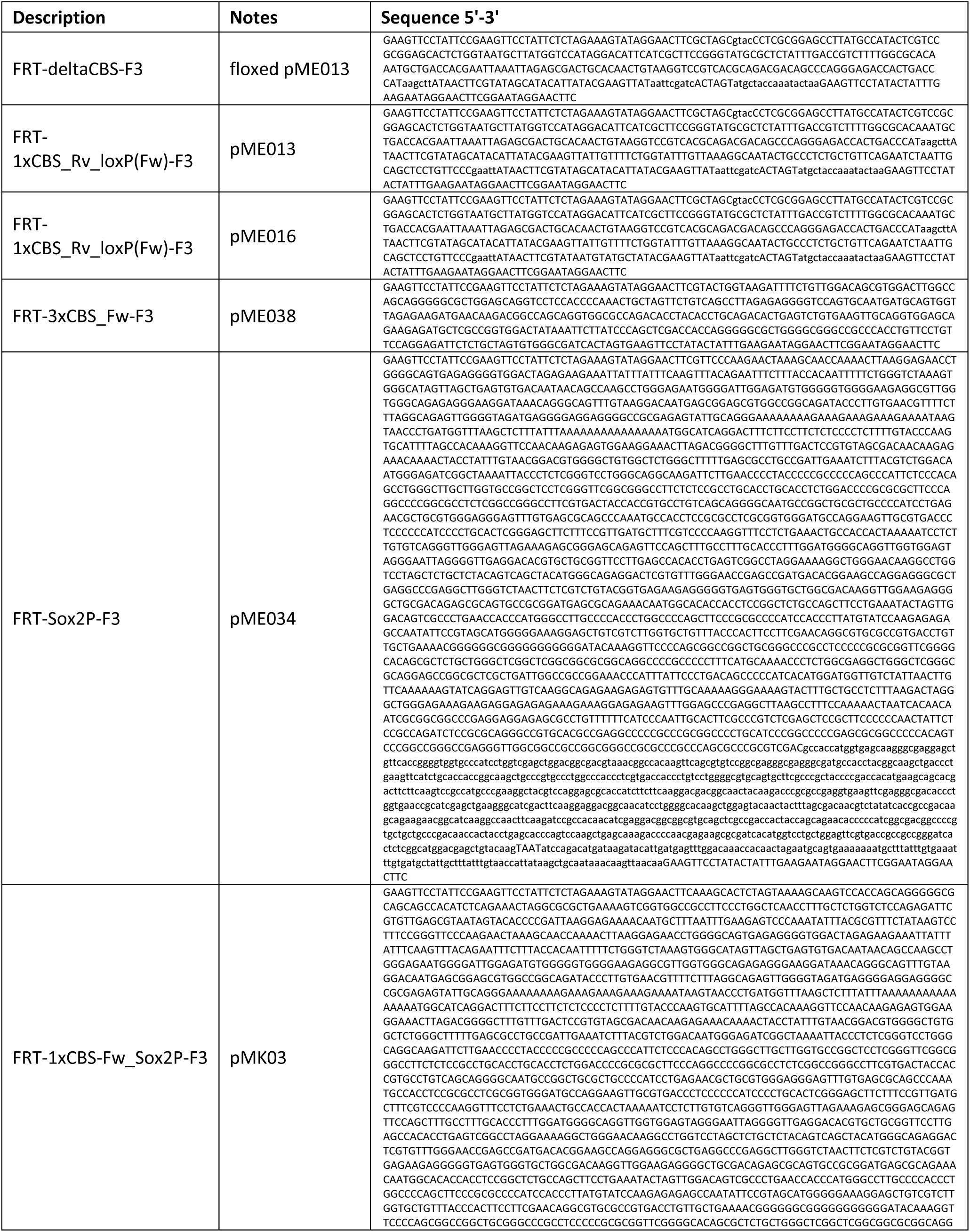
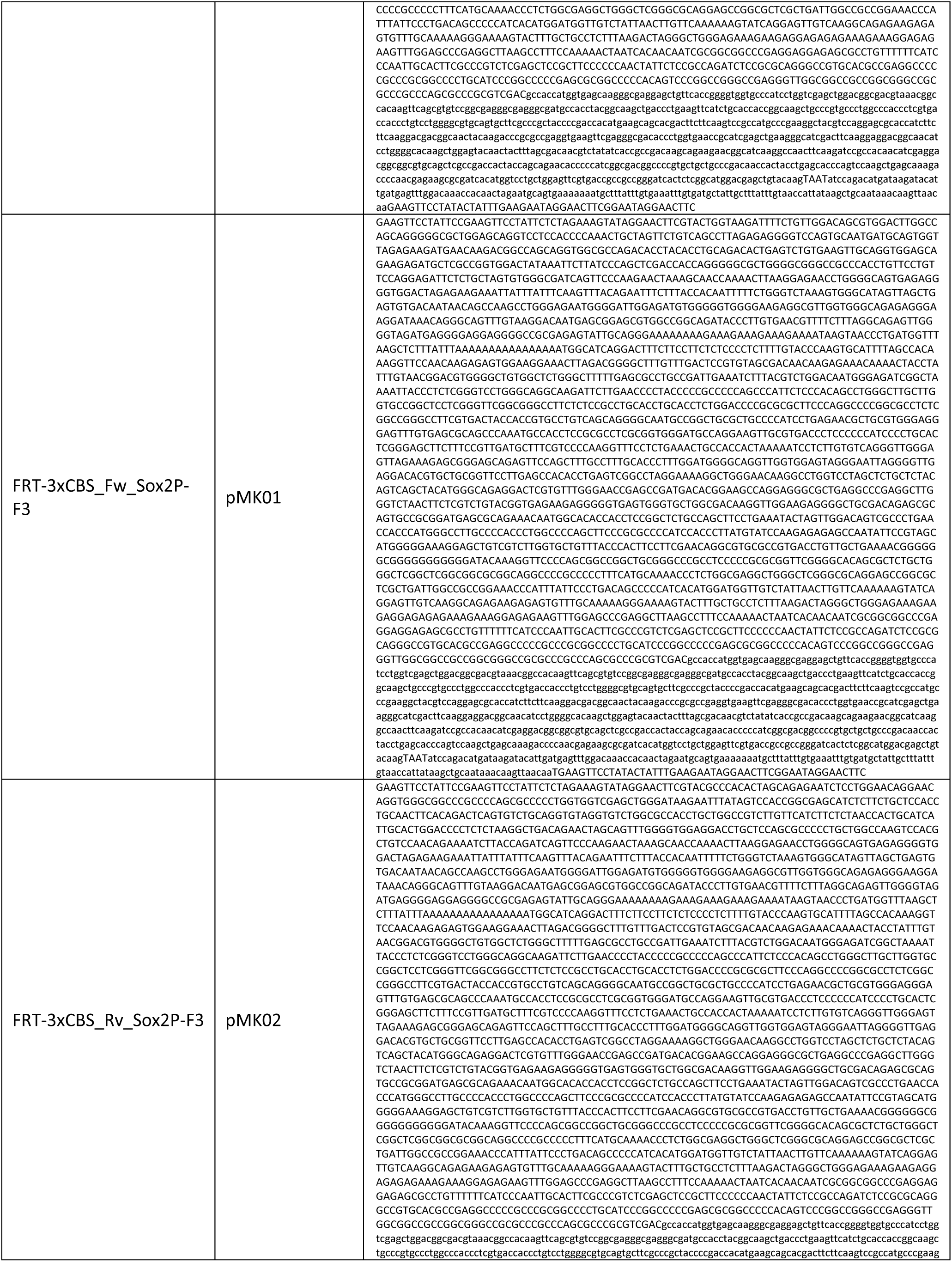
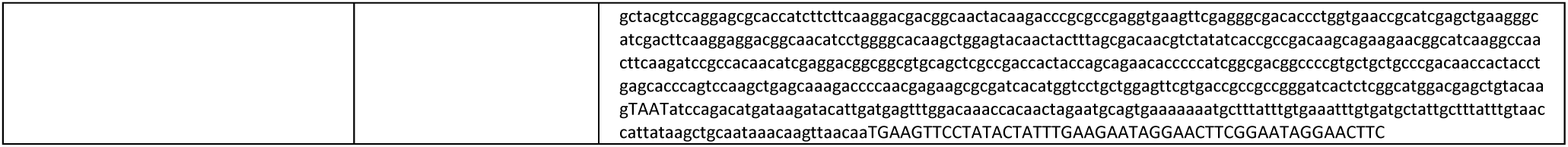
List of sequences used for RMCE.

**Table S6.**
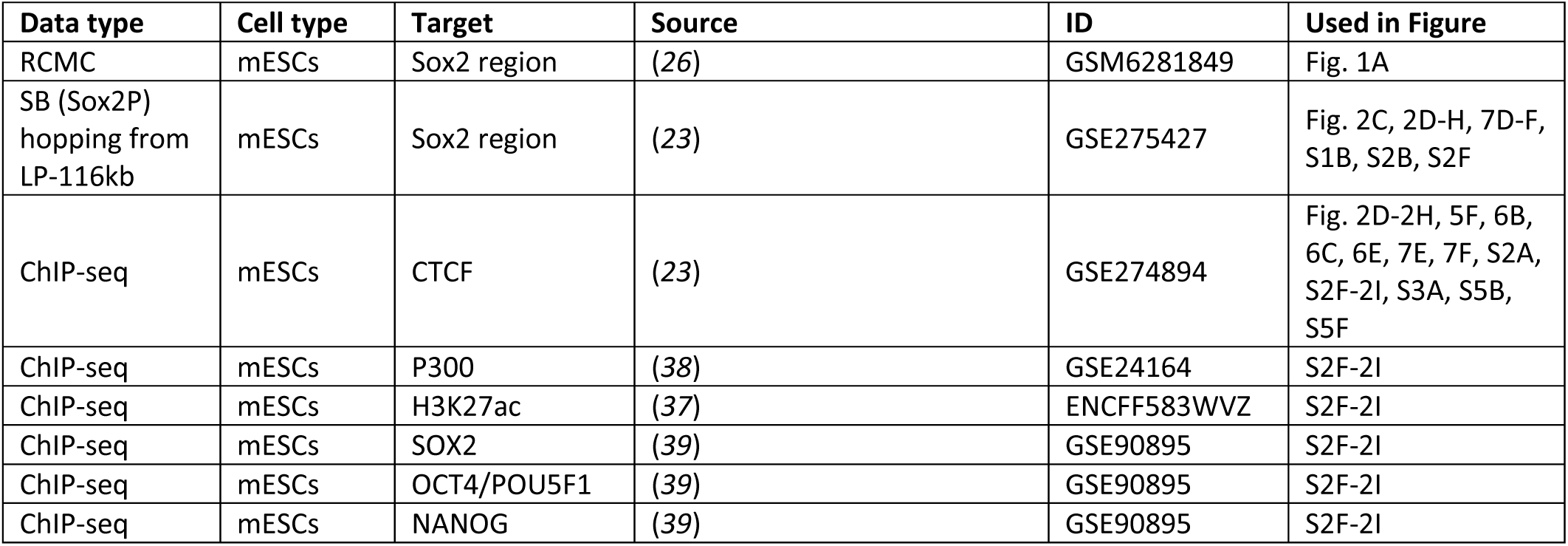
List of external datasets used in this study.

